# Alterations of genomic imprinting appear during the reprogramming of adult neural stem cells

**DOI:** 10.1101/2024.01.23.576820

**Authors:** Raquel Montalbán-Loro, Anna Lozano-Ureña, Laura Lázaro-Carot, Esteban Jiménez-Villalba, Jordi Planells, Adela Lleches-Padilla, Mitsu Ito, Elisabeth J. Radford, Sacri R. Ferrón

**Author notes:** These authors contributed equally to this work (Raquel Montalbán-Loro, Anna Lozano-Ureña, Laura Lázaro-Carot, Esteban Jiménez-Villalba). To whom all correspondence should be addressed, at:* Sacri R. Ferrón, ERI BiotecMed/Departamento de Biología Celular Universidad de Valencia, 46100 Burjassot, Spain.

## Abstract

Genomic imprinting is an epigenetic mechanism that causes monoallelic expression of genes depending on their parental origin. Loss of imprinting (LOI) is associated with cancer progression and human imprinting disorders (IDs), impacting foetal development, metabolism and cognition. Imprinted genes, organized in clusters, rely on methylation at imprint control regions (ICRs), which are differentially methylated regions (DMRs) on both parental chromosomes. Somatic cell reprogramming into induced pluripotent stem cells (iPSCs) is a valuable tool to understand the mechanisms associated with pluripotency and holds promise for generating patient-specific stem cells for therapeutical applications to treat different pathologies such as IDs. Here, we conduct genome-wide RNA-seq and MeDIP-seq analysis on mouse iPSCs derived from adult neural stem cells (NSCs). Our findings reveal a comprehensive alteration in iPSCs transcriptome profile, aligning with DNA hypomethylation. This correlation is pivotal in discerning which modifications in genomic imprinting during the reprogramming process represent undesirable epigenetic abnormalities that could compromiise the quality of iPSCs. Simultaneously, it helps identify genuine epigenetic modifications that are inherently linked to pluripotency, thus ensuring a clearer understanding of the factors influencing iPSC quality and pluripotent potential.

## Introduction

Reprogramming of somatic cells is a valuable tool to understand the mechanisms associated with pluripotency and further opens up the possibility of generating personalised pluripotent stem cells, which provides tremendous implications in regenerative medicine. So far, a variety of cell types have been reprogrammed into induced pluripotent stem cells (iPSCs) including fibroblasts (Takahashi and Yamanaka, 2006), neural progenitor cells (Kim et al., 2009), hepatocytes and gastric epithelial cells (Aoi et al., 2008), B cells (Hanna et al., 2008), pancreatic β cells (Stadtfeld et al., 2008), melanocytes (Utikal et al., 2009) and keratinocytes (Aasen et al., 2008). More concretely, different combinations of reprogramming factors have been used to convert postnatal neural stem cells (NSCs) into iPSCs with similar efficiency and comparable expression profiles to embryonic stem cells (ESCs) (Eminli et al., 2008; Kim et al., 2008). Cell reprogramming involves changes in the transcriptome and chromatin state of the reprogrammed cells to that of a pluripotent stem cell (Montalbán-Loro, 2015). During the acquisition of a pluripotency state, reprogramming factors interfere with methylation by binding to specific promoters or enhancer regions leading to demethylation and activation of the pluripotency genes (Hochedlinger and Jaenisch, 2015), which seems to be crucial for faithful reprogramming (Apostolou et al., 2013).

During mammalian development, the vast majority of genes are expressed or repressed from both alleles. However, there are a small number of genes, termed “*imprinted genes*” that are expressed monoalellicaly from either the maternally or the paternally inherited chromosomes (Ferguson-Smith, 2011). Approximately 200 imprinted genes have been described in mammals and are generally organized in clusters, although examples of singleton imprinted genes do exist (Barlow and Bartolomei, 2014; Ferguson-Smith, 2011; Lassi and Tucci, 2019). An imprinting cluster is usually under the control of a DNA element, called the imprinting control region (ICR) that consists of differentially DNA methylated regions (DMRs) on the two parental chromosomes (Edwards and Ferguson-Smith, 2007; Ferguson-Smith, 2011). ICRs can be divided into those which are methylated on the paternally inherited copy located in distinct intergenic regions, and those with maternally inherited methylation that are frequently located at the promoters (SanMiguel and Bartolomei, 2018; Tucci et al., 2019). Importantly, deletion of an ICR results in loss of imprinting (LOI) of multiple genes in the cluster and this has been associated to several human pathologies and imprinting disorders such as Prader Willi Syndrome (PWS) or Angelman Syndrome (Barlow and Bartolomei, 2014; Ferguson-Smith, 2011; Lassi and Tucci, 2019). Parental specific marks at ICRs are established in the developing germline in an allele-specific manner depending on whether the developing gamete is in the male or female germline (Bartolomei and Ferguson-Smith, 2011; Edwards and Ferguson-Smith, 2007; Ferguson-Smith, 2011). After fertilization, a rapid and extensive reprogramming of the parentally inherited genomes occurs, and most DNA methylation is lost (SanMiguel and Bartolomei, 2018; Smallwood and Kelsey, 2012)(Smallwood and Kelsey, 2012). However, the parental-specific imprints are maintained during this period and a memory of parental origin is propagated into daughter cells during somatic cell divisions (Takahashi et al., 2015).

Genomic imprinting has been found to be variably lost during iPSCs reprogramming, with some imprinted regions more severely affected than others (Perrera and Martello, 2019; Takikawa et al., 2013). Indeed, previous studies have described epigenetic alterations of genomic imprinting during the induction of pluripotency from somatic cells both in mouse and human (Arez et al., 2022; Kim et al., 2013; Lee et al., 2016; Liu et al., 2010; Yagi et al., 2019). For example, the *Dlk1-Dio3* imprinted region has been shown to be hypermethylated in iPSCs causing the altered expression of several genes within the cluster (Liu et al., 2010; Stadtfeld et al., 2008; Stadtfeld et al., 2010). By contrast, other studies have described a global hypomethylation of ICRs during reprogramming, although de novo methylation at a late stage of the reprogramming process has also been reported (Yagi et al., 2019). More recently, it has been suggested that imprinting defects in iPSCs are dependent on the sex of donor cells and on culture conditions and these defects are never rescued upon differentiation (Arez et al., 2022; Bar et al., 2017; Nazor et al., 2012). However, most of these studies examined the expression of a limited number of imprinted genes in iPSCs together with a limited number of ICRs, only providing a partial view of alterations of genomic imprinting in iPSCs. Moreover, previous data from our lab, showed that selective absence of genomic imprinting can occur in physiological conditions in adult NSCs (Ferron et al.2011; 2015). The resultant biallelic expression of imprinted genes is required for normal neurogenesis in the adult brain (Ferrón et al., 2011; Ferrón et al., 2015; Montalbán-Loro et al., 2021) suggesting that these pre-existing alterations in somatic cells are heritable changes in genomic imprinting that might be reversible and context-dependent and are likely to be essential to control stem cell potentiality.

Here we present a global unbiased analysis of impact of reprogramming on adult NSC-derived iPSCs using only the two reprogramming factors *Klf4* and *Oct4*. A genome-wide RNA-seq and MedIP-seq analysis has shown an altered transcriptome profile in iPSCs together with the hypomethylation of their DNA. The study of methylation at all ICRs described in mouse has revealed that in iPSCs the most affected ICRs are the maternally methylated DMRs. These changes in methylation clearly correlated with transcriptional changes of genes located at the affected imprinted clusters. These findings are crucial to understand which alterations of genomic imprinting are undesirable epigenetic abnormalities that might be avoided to preserve cell proliferation and pluripotency quality of iPSCs and which alterations are epigenetic modifications associated to pluripotency. Importantly, these findings may have implications for the therapeutical applications of iPSCs.

## Results

### NSCs from the adult SVZ convert into a pluripotent state with the transduction of *Oct4* and *Klf4*

Previous studies have reported that neurosphere cultures obtained from postnatal day 5 mouse brain endogenously express *Sox2*, *c-myc* and *Klf4* and so can be reprogrammed with *Oct4* alone, or *Oct4* and *Klf4* at a similar efficiency to the reprogramming rate of murine fibroblast with the original four factors (Kim et al., 2008; Kim et al., 2009). To confirm the levels of expression of these transcription factors in NSCs derived from the adult subventricular zone (SVZ), we analysed by quantitative PCR (qPCR) and immunocytochemistry the expression of different neural and pluripotency markers and found that NSCs that consistently expressed neural genes such as *Pax6* or *Olig2*, also expressed significant levels of *Sox2*, *Klf4*, and *c-myc* (**Fig. 1a**). We also confirmed that other genes associated with pluripotency, such as *Oct4*, *Nanog* or *Zfp42* were not expressed in adult NSCs (**Fig. 1a**).

**Figure 1.**
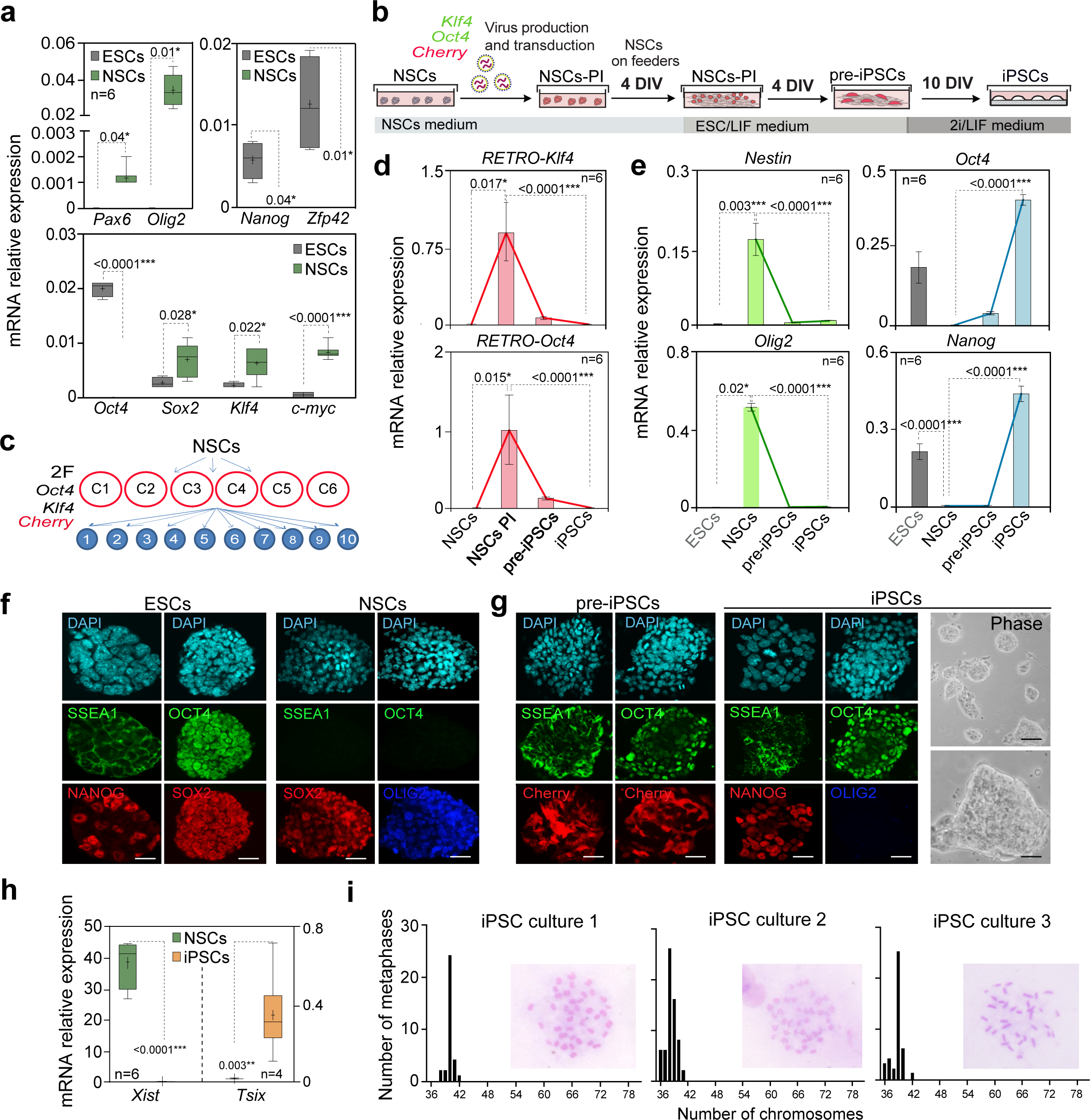
NSCs from the adult SVZ are reprogrammed into iPSCs by exogenously expression of *Oct4* and *Klf4*. **(a)** Quantitative PCR (qPCR) of the neural genes *Pax6* and *Olig2* in adult NSCs and in ESCs pluripotent cultures. The expression of the reprogramming transcription factors *Oct4*, *Sox2*, *Klf4* and *c-myc*; and the pluripotency markers *Nanog* and *Zfp42*, were also analysed in adult NSCs and ESCs. **(b)** Schematic of the reprogramming protocol: Plat-E transfection with plasmids expressing *Oct4*, *Klf4* and *Cherry* was used to generate viral particles to infect NSCs 3 days after. Infected NSCs (NSCs-PI) were seeded on *feeders* 4 days after infection. *Cherry* and SSEA-1 positive clone-likes aggregates appeared (pre-iPSCs) after 4 days and the culture medium was then changed to 2i/LIF medium. iPSCs appeared 10 days after changing the medium. **(c)** Schematic representing the performed reprogramming assay. NSCs isolated from the adult SVZ of six different mice (culture 1 to 6) were transduced with *Oct4* and *Klf4* (2F: 2-factors). Once iPSCs were generated, ten clones of each culture were picked and expanded for further characterization. **(d)** qPCR of *Klf4* and *Oct4* retroviral expression in adult NSCs, NSCs-PI, pre-iPSCs and iPSCs**. (e)** qPCR of the neural genes *Nestin* and *Olig2* in the same samples (left panel). qPCR of the pluripotency-related genes *Oct4* and *Nanog* in adult NSCs, pre-iPSCs and iPSCs. ESCs were used as a positive control. **(f)** Immunocytochemistry images for SSEA-1 and OCT4 (green; middle panel) and for NANOG and SOX2 (red, lower panel) in ESCs and NSCs. Immunocytochemistry for OLIG2 (blue; lower panel) in NSCs is also shown. **(g)** Immunocytochemistry images for SSEA-1 and OCT4 (green; middle panel) and for Cherry (red, lower panel) in pre-iPSCs and iPSCs. Immunocytochemistry for OLIG2 (blue; lower panel) and NANOG (red; lower panel) in iPSCs are also shown. Phase contrast images of the iPSCs generated (right panel). **(h)** qPCR of *Xist* and *Tsix* genes in NSCs and female iPSCs. **(i)** Quantification of the number of chromosomes per metaphase in the three selected clones from the iPSCs cultures. Normal chromosomic dotation was observed (purple bars). Examples of phase contrast images for Leishman staining are included. At least 50 metaphases were counted for each iPSCs line. *Gapdh* was used to normalize gene expression data. DAPI was used to counterstain nuclei. All error bars show s.e.m. P-values and number of samples are indicated. Box and whisker plots show the mean (+), median (horizontal line in box) and maximum and minimum values (whiskers). Scale bars in f and g: 20 µm (phase images in g: 40 µm in upper panel and 5 µm in lower panel).

We later developed a reprogramming protocol using retrovirus by transfecting Plat-E packing cells with retroviral plasmids expressing the transcription factors *Oct4* and *Klf4*. A retrovirus for *Cherry* was also used as a reporter of the exogenous expression of the reprogramming factors (**Fig. 1b**). After two days of retrovirus production, a combination of supernatants with retrovirus for *Oct4* and *Klf4* were used to co-transduce adult NSCs (2 factors condition, 2F) in combination with a retrovirus for the reporter Cherry (**Fig. 1b**). NSCs infected with both *Oct4* and *Klf4* transcription factors were grown on a feeder layer of embryonic fibroblasts with the cytokine leukaemia inhibitory factor (LIF) (**Fig. 1b**). Notably, cultures started to form clone-like aggregates 10 days after infection (**Fig. S1a**). These aggregates were large, with poorly defined edges and some of them expressed the pluripotency marker Stage-specific embryonic antigen 1 (SSEA-1) (**Fig. S1a**). Retroviral vectors are transcriptionally silent in pluripotent stem cells (Hotta and Ellis, 2008). However, most of the clones formed were still positive for Cherry, indicating that despite expressing SSEA-1 they were only partially reprogrammed (**Fig. 1b** and **S1a**). We considered this state of cells as pre-iPSCs.

To promote a ground state of pluripotency of these pre-iPSCs, we applied molecularly defined conditions by neutralizing inductive differentiation stimuli combining molecule inhibitors for dual inhibition (2i) of mitogen-activated protein kinase signalling (MEKi) and glycogen synthase kinase-3 (GSK3) (Silva et al., 2008). In this serum-free culture medium, LIF was also added, which maximized clonogenic self-renewal of pluripotent cells (**Fig. 1b**) (Ying et al., 2008). This protocol of reprogramming was repeated in six independent cultures of adult NSCs (**Fig. 1c**). To further determine the pluripotent state of the clones generated, (Hotta and Ellis, 2008)we next studied the acquisition of pluripotency markers and silencing of neural genes, together with the silencing of retroviral vectors,(**Fig. 1d-g**). After 10 days in the new controlled 2i/LIF conditions, we obtained ESC cell-like colonies that did not express Cherry (**Fig. S1b**), suggesting that the cells had been fully reprogrammed into iPSCs (**Fig. 1g** and **S1b**).

Consistent with the cell cultures, expression studies performed by qPCR corroborated that post-infected NSCs (NSCs PI) expressed high levels of both *Oct4* and *Klf4* retroviral transgenes and that although lower, they were still present in pre-iPSCs (**Fig. 1d**). Moreover, although expression levels of the neural markers *Nestin* and *Olig2* were already downregulated (**Fig. 1e**), pre-iPSCs only showed a slight increase in endogenous *Oct4* levels (**Fig. 1e**) whereas the expression of other pluripotency markers such as *Nanog* and *Zfp42 (Rex1)* was undetectable in pre-iPSCs (**Fig. 1e** and **S1c**). These results confirmed that pre-iPSCs were an intermediate state in which critical attributes of true pluripotency, including stable expression of endogenous *Oct4* and *Nanog,* were not attained yet. However, complete downregulation of retroviral transgenes, essential for full reprogramming, was corroborated in derived iPSCs compared to the original infected NSCs (**Fig. 1d**). This was accompanied by the stable induction of endogenous pluripotency-related genes *Oct4, Nanog* and *Zfp42* (**Fig. 1e** and **S1c**). Importantly, iPSCs and ESCs expressed similar levels of these pluripotency markers (**Fig. 1e** and **S1c**) consistent with the acquisition of a pluripotent state. Moreover, the expression of neural-specific genes in NSCs, *Nestin* and *Olig2*, was absent in iPSCs (**Fig. 1e**). Immunocytochemistry studies confirmed the presence of Cherry in pre-iPSCs but not in iPSCs (**Fig. 1f** and **S1b**). Finally, immunocytochemistry for OCT4 and NANOG and the histochemistry for the Alkaline phosphatase confirmed the pluripotency of iPSCs generated (**Fig. 1g** and **S1d**).

We next corroborated the naïve pluripotency of iPSCs by evalulating the reactivation of the inactive X chromosome in female iPSCs (Cantone and Fisher, 2017; Ohhata and Wutz, 2013; Pasque and Plath, 2015). To ascertain the X chromosome status in iPSCs derived from NSCs, a qPCR of *Xist*, responsible for X chromosome inactivation and *Tsix*, repressor of *Xist*, was performed (Cantone and Fisher, 2017; Janiszewski et al., 2019). This study revealed a reduction of *Xist* expression and an increase of the expression of *Tsix* in iPSCs, resembling closely the patterns observed in ESCs (**Fig. 1h**). These findings were coincident with X chromosome reactivation, confirming the acquisition of a fully pluripotent state in iPSCs.

Genetic variations, such as aneuploidy or polyploidy, may be introduced during the generation of iPSCs (Liu et al., 2020; Vaz et al., 2021). Hence, we next conducted a karyotype analysis of the generated iPSCs. Ten clones of iPSCs were isolated and expanded from each culture, and three were chosen based on their gene expression profile (displaying high expression of pluripotency genes and low expression of neural and retroviral genes) to evaluate their genomic stability. The analysis revealed that the vast majority of the lines analysed (93%) exhibited a normal karyotype with approximately 40 chromosomes per metaphase (**Fig. 1i**). iPSCs with chromosomal abnormalities were excluded from further studies.

### iPSCs derived from adult NSCs differentiate *in vitro* and *in vivo* into cells from the three gem layers

Pluripotent stem cells have the potential to differentiate into cells of the three germ layers, endoderm, mesoderm and ectoderm. This differentiation potential is typically confirmed by demonstrating the capacity of the iPSCs to form three-dimensional structures called embryoid bodies (EBs), comprising cells characteristic of these germ layers (Höpfl et al., 2004). EBs emulate the structure of developing embryos and serve as a model to obtain various cell lineages (Spelke et al., 2010). Thus, to explore the developmental capacity of iPSCs, we induced the formation of EBs by subjecting cells to conditions that are adverse to pluripotency and proliferation using the hanging drop method (**Fig. 2a**). Suspended iPSCs on the dish lid aggregate at the base of the drop (**Fig. 2a**), consistently generating uniform EBs (Höpfl et al., 2004). ESC differentiation was utilized as a comparative control (**Fig. S2**).

**Figure 2.**
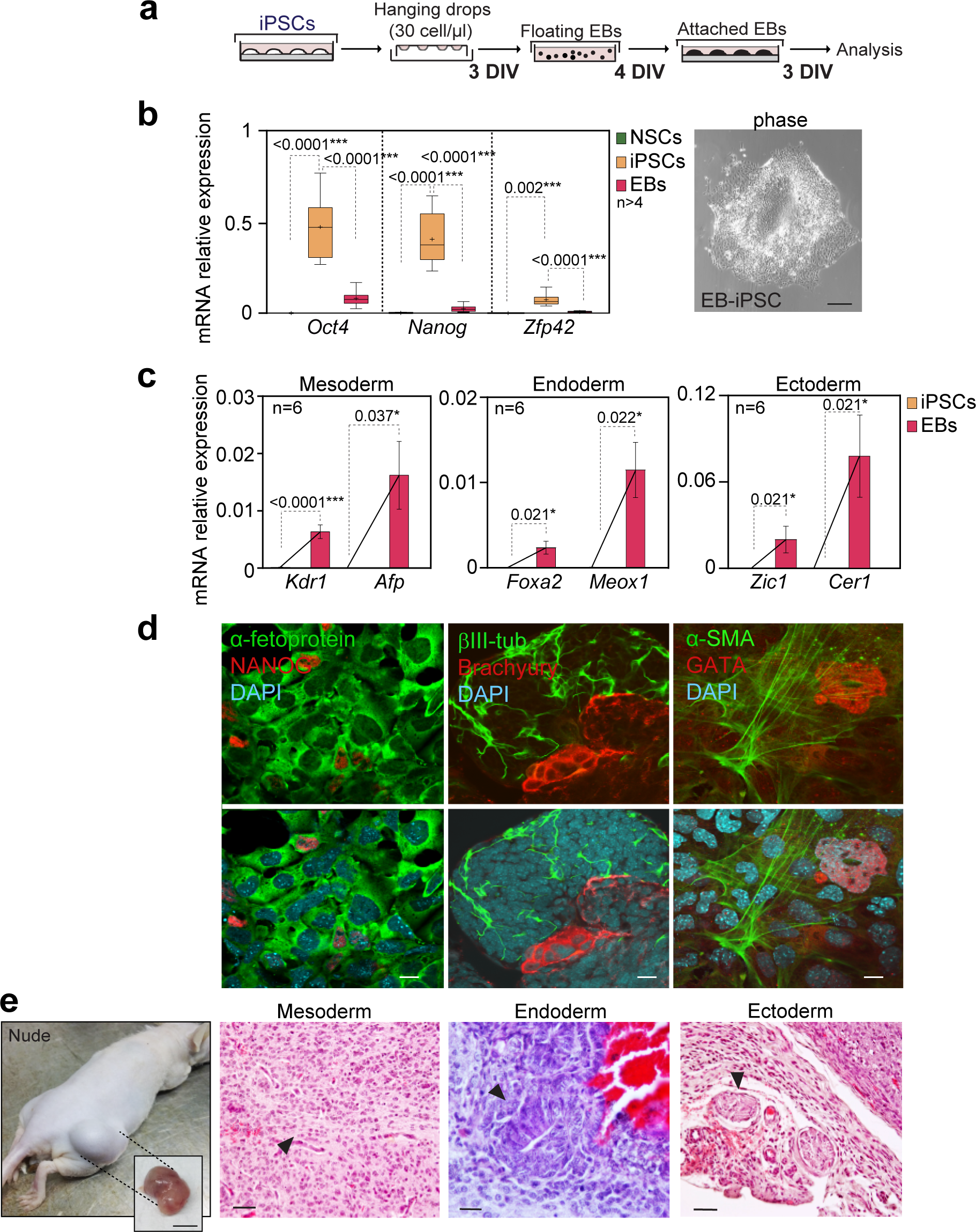
iPSCs generated from NSCs are able to differentiate into cells of the three germ layers *in vitro* and *in vivo*. **(a)** Schematic representation of the embryoid bodies (EB) assay using the “*hanging drops*” method. iPSCs were dissociated and the cell suspension was distributed in drops in a plate that was incubated upside-down for 3 days. Then, the plate was inverted and culture media was added. Incipient EBs were incubated 4 days more. Finally, EBs were seeded in gelatine pre-treated to allow differentiation. After 3 days, samples were analysed**. (b)** qPCR of *Oct4*, *Nanog* and *Zfp42* genes in NSCs, iPSCs and EBs derived from iPSCs (left panel). A phase contrast image of the EBs is included (right panel). **(c)** qPCR of *Kdr1* and *Afp* (mesoderm), *Foxa2* and *Meox1* (endoderm) and *Zic1* and *Cer1* (ectoderm) genes in NSCs, iPSCs and EBs derived from iPSCs. **(d)** Immunocytochemistry images for α-fetoprotein (green) and NANOG (red) (left panel), for βIII-tubulin (green) and Brachyury (red) (middle panel) and for α-SMA (green) and GATA (red) (right panels) in EBs derived from iPSCs. **(e)** Image of a teratoma developed in the dorsolateral area of immunocompromised *Nude* mice 2 weeks after the injection. A detailed image of the teratoma after its extraction is included (left panel). Histological analysis of teratomas using haematoxylin-eosin dyes (right panel). Epithelial cells derived from ectoderm, muscle fibres derived from mesoderm and columnar epithelium from endoderm are shown and indicated with arrowheads. DAPI was used to counterstain nuclei. *Gapdh* was used to normalize gene expression data. All error bars show s.e.m. of at least 6 samples. Box and whisker plots show the mean (+), median (horizontal line in box) and maximum and minimum values (whiskers). Scale bar in b: 10 µm; in d: 50 µm; in e: 1 cm (left panel) and 20 µm (right panel).

Initially, we determined the gene expression in iPSCs-derived EBs using RT-PCR analysis. The results indicated a downregulation of the pluripotency markers *Oct4*, *Nanog* and *Zfp42* after differentiation of iPSCs (**Fig. 2b**). Additionally, a significant expression for the mesoderm markers *Kdr1* and *Afp*, the endoderm markers *Foxa2* and *Meox1,* and the ectoderm markers *Zic1* and *Cer1,* were observed, confirming the presence of cells representing the three germ layers within the generated EBs (**Fig. 2c)**. Immunocytochemistry utilizing specific antibodies for α-fetoprotein and GATA4 (Endoderm), βIII-tubulin (neuroectoderm) and, Brachyury and α-SMA (mesoderm) further corroborated these results (**Fig. 2d** and **S2b**).

We next evaluated the in vivo pluripotent capacity of iPSCs derived from adult NSCs using the teratoma assay. Teratomas are non-malignant tumours that result from uncontrolled expansion and disorganized differentiation of pluripotent cells. When ESCs are transplanted into immunocompromised Nude mice, they trigger teratoma formation (Prokhorova et al., 2009; Przyborski, 2005). In this study, iPSCs were injected into the dorsolateral area in the subcutaneous space of immunocompromised *Nude* mice. Within 10 days, teratomas became visible to the naked eye, reaching approximately 2 cm of diameter 20 days after the injection (**Fig. 2e**). Subsequent histological analysis of the teratomas upon sacrifice revealed disorganized tumoral cytoarchitecture, while staining with haematoxylin-eosin confirmed the presence of cells representing all three germ layers was disorganized and the presence of cells from all three germ layers was confirmed by haematoxylin-eosin staining (**Fig. 2e**). These teratoma cells demonstrated differentiation into ectodemal secretory epithelium, mesodermal cartilage and endodermal gut epithelium derivatives (**Fig. 2e).** These finding together confirmed the *in vitro* and *in vivo* pluripotency of the iPSCs derived from adult NSCs.

### The global transcriptome profile is modified in iPSCs reprogrammed from adult NSCs

RNA-seq analysis was performed to investigate the transcriptional differences between NSCs and iPSCs (GEO Accession# XXXXXX). Principal component analysis (PCA) conducted on the RNA-seq data distinctly separated iPSCs from NSCs (**Fig. 3a**). Comparison of NSCs versus iPSCs unveiled 11,363 differentially expressed genes, representing 50% of all genes expressed in NSCs (**Fig. S3a**). Among these alterations, 6,062 genes were downregulated while 5,301 were upregulated in iPSCs compared to NSCs (**Fig. 3b** and **S3a**). As expected, RNA-seq confirmed the downregulation of neural-specific genes like *Olig2, Nestin* and *Zic1,* alongside the upregulation of pivotal pluripotency-associated genes involved in pluripotency such as *Oct4, Nanog* and *Zfp42* in iPSCs compared to NSCs (**Fig. 3c**). Based on RNA-seq data a “*Gene Set Enrichment Analysis*” (GSEA) was performed (**Fig. S3b**). Remarkably, GSEA identified 39 gene sets exhibiting downregulation in iPSCs encompassing important biological functions including focal adhesion, axon guidance and apoptosis (**Fig. S3b**). Conversely, 97 gene sets showed upregulation, involving pathways associated with cancer and MAPK signalling, among others (**Fig. S3b**). The Gene Ontology (GO) analysis of the altered genes between NSCs and iPSCs revealed that reprogramming significantly affected the biological process of monosaccharide catabolism, while the cellular component most influenced by differentially expressed genes was identified as the microtubule end (**Fig. S3c**).

**Figure 3.**
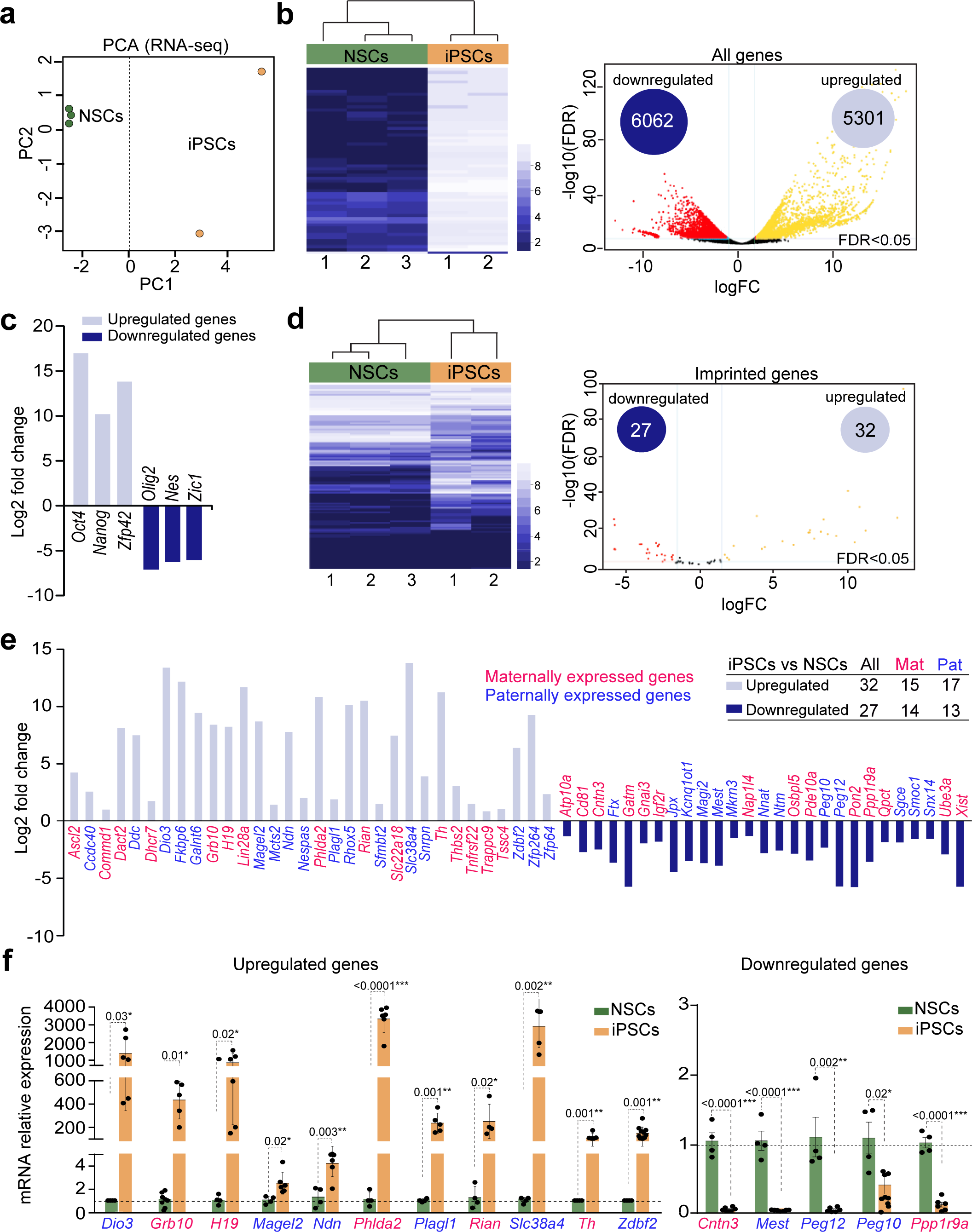
Expression of imprinted genes in adult NSCs is importantly regulated during the reprogramming process. **(a)** Principal component analysis (PCA) from RNA-seq of 3 NSCs and 2 iPSCs cultures where samples are represented based on their total expression levels. **(b)** Heatmap representing the 60 most differentially expressed genes between adult NSCs and iPSCs (FDR<2.5e-68). Upregulated genes are shown in light blue, whereas downregulated genes are shown in dark blue (left panel). Volcano plot for all genes differentially expressed based on RNA-seq data (right panel). The number of downregulated and upregulated genes in iPSCs compared to NSCs are indicated. **(c)** Log_2_ fold change from the RNA-seq data of the three pluripotency genes *Oct4, Nanog* and *Zfp42* and of the three neural genes *Olig2, Nestin (Nes)* and *Zic1*. **(d)** Heatmap representing the expression of imprinted genes in NSCs and iPSCs. Non-expressed imprinted genes were discarded from subsequent analysis (left panel). Volcano plot for differentially expressed imprinted genes based on the RNA-seq data (right panel). **(e)** Graph showing the Log_2_ fold change of upregulated (light blue) and downregulated (dark blue) imprinted genes in iPSCs compared to NSCs. Number of maternally (pink) and paternally (blue) expressed genes that are altered is included. **(f)** RT-PCR validating the levels of expression of several imprinted genes in NSCs and iPSCs. *Gapdh* was used to normalize gene expression data. All error bars show s.e.m. P-values and number of samples are indicated.

### Expression of imprinted genes is altered during the reprogramming of NSCs into iPSCs

The majority of imprinted genes are expressed in the brain and recent evidence suggests that genomic imprinting can be selectively lost in particular cell types or at specific developmental time points (Ferrón et al., 2011; Ferrón et al., 2015; Kim et al., 2013; Montalbán-Loro et al., 2021; Tucci et al., 2019). Moreover, some alterations of genomic imprinting have been observed during the reprogramming of somatic cells (Perrera and Martello, 2019; Takikawa et al., 2013).These changes have an impact on stem cell plasticity suggesting that genomic imprinting may be a mechanism employed to modulate gene dosage to control stem cell potential (Ferrón et al., 2011; Perez et al., 2016). Therefore, we next focused on the study of the regulation of imprinted genes during the reprogramming process by using the RNA-seq analysis performed in NSCs and iPSCs. RNA-seq data identified 59 imprinted genes that were differentially expressed (FDR<0.05) between iPSCs and NSCs, representing around 50% of all imprinted genes initially expressed in NSCs (**Fig. 3d,e** and **S3a**). Among them, we observed similar changes in both paternally and maternally expressed genes (**Fig. 3e**). A validation of the expression of most of these imprinted genes was performed by qPCR in NSCs and iPSCs (**Fig. 3f** and **Fig. S4a**) confirming that the acquisition of a pluripotent state involves significant transcriptional changes also of imprinted genes.

### A global hypomethylation of DNA was observed in iPSCs generated from NSCs

Previous studies have shown that changes in DNA methylation patterns are essential for successful cell reprogramming, exemplified by the necessity for methylation loss at the promoter of pluripotency genes (Hochedlinger and Jaenisch, 2015; Lee et al., 2014; Takahashi and Yamanaka, 2006). Moreover, DNA methylation represents one of several epigenetic mechanisms employed by cells to regulate gene expression during cell fate decisions (Parry et al., 2021). In order to characterise methylation changes genome-wide, a MeDIP-seq analysis was performed in iPSCs and NSCs. This experimental approach uses a 5-methylcytosine (5mC) antibody to enrich for DNA fragments containing this modification followed by high throughput sequencing. A principal component analysis of MeDIP-seq data showed clear segregation between the NSCs of origin and the iPSCs generated (**Fig. 4a**).

**Figure 4.**
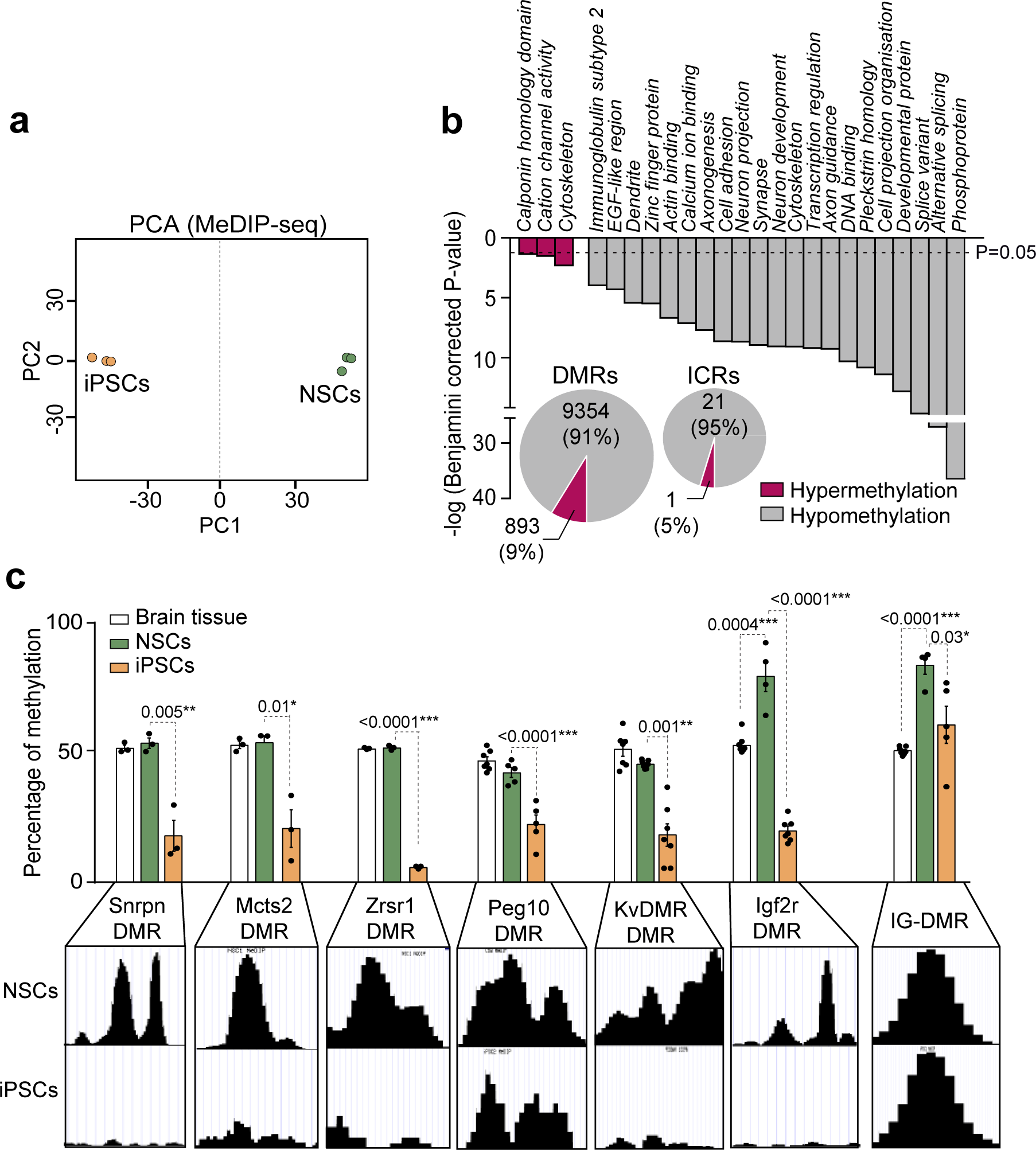
A global hypomethylation of DMRs is observed in iPSCs compared to adult NSCs. **(a)** Principal component analysis (PCA) from MeDIP-seq of 3 NSCs and 3 iPSCs cultures based on their global methylation levels. **(b)** *Gene Set Enrichment Analysis* (GSEA) showing statistically significant differences between NSCs and derived iPSCs. Several sets of genes involved in different biological functions showed changes in methylation. Total number of differentially methylated regions (DMRs) that show hypomethylation (gray) or hypermethylation (pink) in the whole genome of iPSCs. The number of imprinting control regions (ICRs) that are altered in iPSCs compared to NSCs is also shown. **(c)** Quantification by pyrosequencing of the percentage of methylation at different DMRs located at ICRs in NSCs (green) and iPSCs (orange). The maternally methylated DMRs (pink), *Snrpn* DMR, *Mcts*2 DMR, *Zrsr1* DMR, *Igf2r* DMR, *Peg10* DMR and KvDMR showed hypomethylation, whereas the paternally methylated IG-DMR (blue) was not altered in iPSCs compared to NSCs. All error bars show s.e.m. P-values and number of samples are indicated.

It has been described that iPSCs have lower levels of methylation than somatic cells, suggesting that demethylation is an important chromatin feature to achieve pluripotency (Lee et al., 2014). In agreement with this, the genome-wide analysis performed demonstrated that 9354 DMRs exhibited reduced methylation levels in iPSCs compared to NSCs, while only 893 DMRs had elevated levels of methylation (**Fig. 4b**). Based on these MeDIP-seq data, we performed a GSEA which identified 21 sets of genes that showed hypomethylation and that were involved in several important biological functions including cell adhesion, cytoskeleton, DNA binding, splice variant, alternative splicing, zinc finger proteins and transcriptional regulation (**Fig. 4b**). Only three sets of genes showed significant hypermethylation levels and these were related to cytoskeleton and cation channel activity (**Fig. 4b**). To establish a link between the altered DNA methylation pattern and transcriptional activity we determined whether changes of expression in pluripotency and neural genes observed in iPSCs compared to NSCs correlated with a gain or loss of the methylation especially at these promoter regions. As expected, low levels of methylation correlated with the upregulation of *Nanog*, *Oct4* and *Zfp42* in iPSCs compared to NSCs (**Fig. S4b**). Consistently, a gain of methylation was found at the promoters of the downregulated genes such as *Nestin*, *Zic1* and *Olig2* (**Fig. S4b**). These data validated the MeDIP-seq protocol and confirmed that the acquisition of a pluripotent state requires reprogramming of methylation state genome-wide, associated with a new gene expression profile in iPSCs.

### Methylation of DMRs at ICRs is modified during the reprogramming of NSCs into iPSCs

Due to the importance of DNA methylation at DMRs to maintain levels and allele-specific patterns of imprinted gene expression, we focused the analysis of methylation on imprinted clusters. Based on MeDIP-seq data, from 33 imprinted DMRs analysed, the methylation profile was substantially modified in 22 of them in iPSCs compared to NSCs (**Fig. 4b** and **5**). Interestingly, 21 of these DMRs (95%) showed hypomethylation and only one DMR (5%) was hypermethylated in iPSCs (**Fig. 4b**). To validate the MeDIP-seq data we used pyrosequencing to quantify methylation levels of DNA subjected to sodium bisulfite treatment. This confirmed that the maternally methylated DMRs, including the *Snrpn, Mcts2*, *Zrsr1*, *Igf2r*, *Peg10* DMRs and KvDMR were hypomethylated, whereas the paternally methylated IG-DMR was not altered in iPSCs compared to NSCs (**Fig. 4c**). These results suggested that the aberrations of DNA methylation at ICRs that occurred during the acquisition of a pluripotent state seem to be more frequent in maternally methylated DMRs and thus the control of the gene expression of imprinted genes at these clusters could be specifically altered in iPSCs.

**Figure 5.**
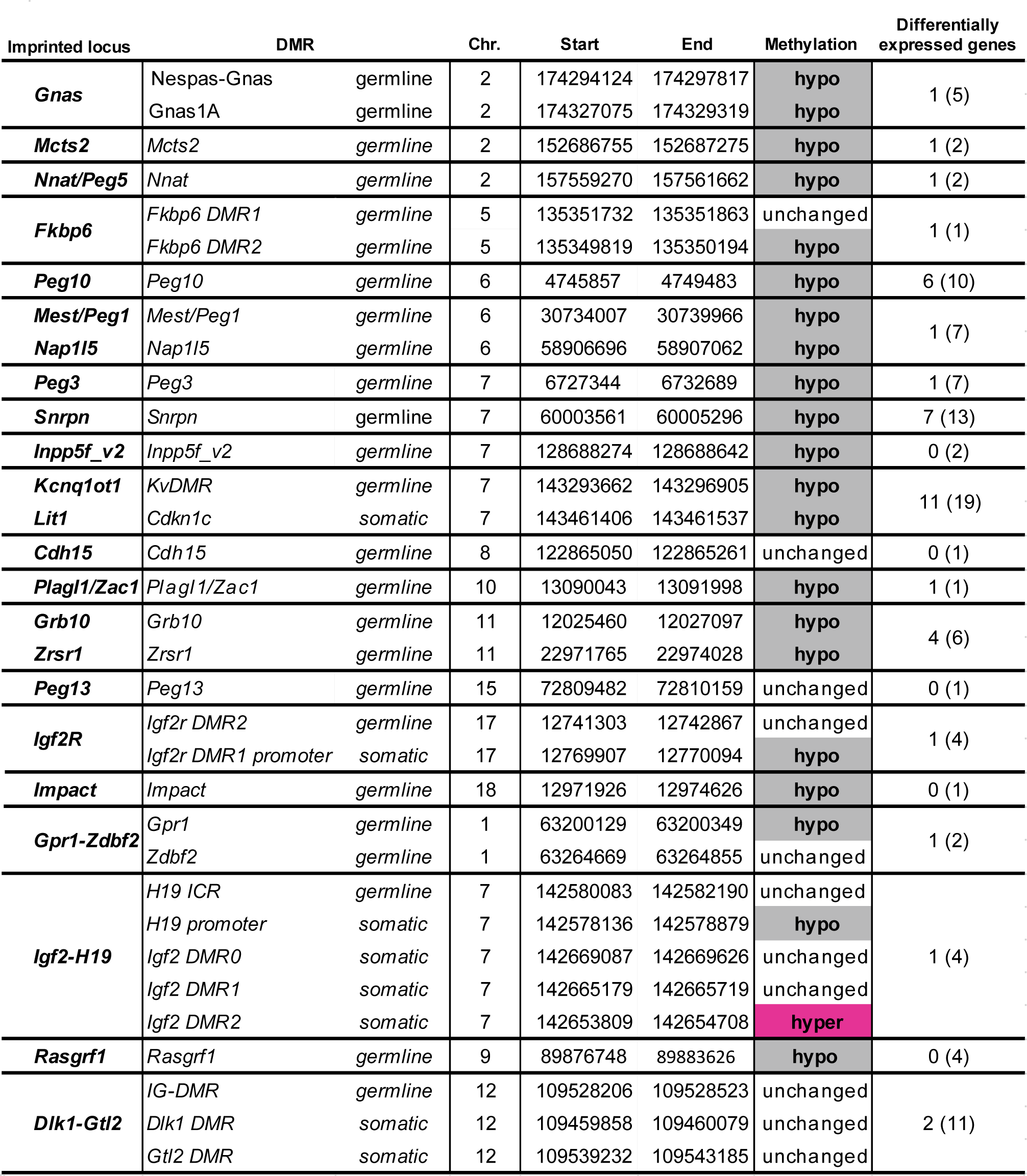
Alterations of methylation and gene expression in different imprinting clusters. The methylation status of DMRs within 23 imprinting cluters is shown. Maternally and paternally methylated DMRs are included. Chromosome (Chr) where the imprinted cluster is located and the number of genes which expression is altered within these clusters are also shown. The start and end nucleotide position based on the Genome Reference Consortium Mouse Build 38 (GRCm38/mm10) are also indicated.

We next studied the relationship between methylation at different ICRs with the expression levels of imprinted genes within the imprinted cluster regulated by these DMRs. Hypomethylation at maternally methylated DMRs correlated with changes in the expression levels of many genes within the clusters (**Fig. 5** and **Fig. S5**). For example, genes from the *Kcnq1ct1*, *Snrpn*, *Grb10* or *Peg10* imprinted clusters were significantly altered (**Fig. 5** and **Fig. S5a**) suggesting that changes in DNA methylation patterns at imprinted loci during reprogramming resulted in changes of gene expression. Maternally methylated DMRs are promoter DMRs, thus our results suggest that reprogramming of NSCs into iPSCs alters this type of DMRs but not intergenic DMRs within imprinting clusters. Indeed, paternally methylated DMRs such as the IG-DMR and *Igf2-H19* DMR, showed no alterations of the levels of expression in most of the genes within these clusters after reprogramming (**Fig. 5** and **Fig. S5b**), however *H19* gene expression, located at the Igf2-H19 paternally methylated cluster was upregulated in iPSCs correlating in this case with loss of methylation at the H19 promoter somatic DMR (**Fig. S5b**).

## Discussion

Somatic cell reprogramming is a very useful tool to understand the mechanisms involved in the acquisition of a pluripotent state and gives information about the differentiation program attributed to the cell of origin. Since the discovery of the reprogramming process in 2006 by Takahashi and Yamanaka (Takahashi and Yamanaka, 2006), researchers have reported the generation of iPSCs from different somatic tissues using the OSKM cocktail. Moreover, several assays have reported a successful reprogramming with a reduced number of factors (Kim et al., 2008; Kim et al., 2009). NSCs significantly express *c-myc*, *Sox2* and *Klf4* suggesting an intermediate state between differentiated and embryonic stem cells (Kim et al., 2008; Kim et al., 2009). The presence of these three factors suggests the possibility to reprogram NSCs using only *Oct4* as exogenous factor (Kim et al., 2009). However, a low efficiency to obtain pluripotent colonies was shown in these studies (Kim et al., 2008; Kim et al., 2009).

Analysis of previously described iPSCs revealed several retroviral integrations for all four factors. Thus, reprogramming of adult NSCs with only one or two factors has important implications as reducing the number of factors decreases the chance of retroviral insertional mutagenesis (Aoi et al., 2008; Wernig et al., 2008). Moreover, *c-myc* is a well-characterised oncogene, and therefore its overexpression may confer a tumorigenic risk (Kuttler and Mai, 2006). Thus, generation of iPSCs lacking *c-myc* represents an important advantage where the ultimate goal is the clinical utilisation of the iPSCs generated. Our study shows that *Oct4* and *Klf4* are sufficient to induce pluripotency in adult NSCs, which demonstrates their crucial role in the process of reprogramming and support the hypothesis that NSCs might represent an intermediate state between differentiated and pluripotent cells. Our study also demonstrates that during the reprogramming process there exist partially reprogrammed cells, called pre-iPSCs that exhibit ESC-like morphology but retain retroviral gene expression and incomplete expression of pluripotency-associated genes, such as *Nanog*. These cells can be easily promoted to full pluripotent state by applying defined conditions using inhibitors for MEK and GSK and in the presence of LIF (2i/LIF), as previously described (Silva et al., 2008). In response to 2i/LIF, pre-iPSCs generated from NSCs rapidly reach a fully pluripotent state with phenotypic and functional characteristic of ESCs. Therefore, *Nanog* occupies a central position in the transcriptional network in the regulation of pluripotency being essential for the formation of iPSCs also from NSCs. In contrast, it has been suggested that *Nanog*, although an important mediator of reprogramming, is not required for establishing pluripotency in murine fibroblasts (Carter et al., 2014), suggesting that different cells of origin may require distinct factors to acquire a pluripotent state.

Although iPSCs exhibit superficial similarity with ESCs in their morphology, gene expression profile and pluripotency, new evidence demonstrates substantial molecular and functional differences among iPSCs derived from distinctive cell types, including the tumorigenic potential and gene expression. This suggests that the somatic cell of origin affects the properties of resultant iPSCs (Polo et al., 2010). For example, iPSCs derived from astrocytes possess more potential for neuronal differentiation compared to fibroblasts-iPSCs (Tian et al., 2011). Therefore, neural-derived iPSCs may retain a “*memory*” of the central nervous system, which confers additional potential (Nashun et al., 2015). Importantly, our results demonstrate that iPSCs obtained from adult NSCs are able to give rise to cells of the three germinal layers after differentiation confirming their pluripotency capability. Strikingly, these iPSCs derived from adult NSCs have a higher differentiation ability to form ectoderm compared to ESCs after differentiation, but also expression of endodermal lineage determinants is induced more efficiently, supporting the hypothesis of iPSCs derived from NSCs having an additional potential compared to non-stem cell derived iPSCs.

The determinants and the temporal order of epigenetic processes leading from a differentiated to a pluripotent cell during iPSC derivation are poorly understood. However, it is clear that to acquire a pluripotent state, cells must erase differentiation-specific epigenetic marks to achieve an ESC-like state implying important transcriptome changes {Hochedlinger, 2015 #62}. Genome-wide expression analysis using next generation sequencing in iPSCs and in the adult NSCs of origin, show that several markers are sequentially activated or repressed after the induction of reprogramming. More concretely and as expected, repression of the neural markers *Nestin* and *Olig2*, correlate with the activation of the pluripotency genes *Oct4*, *Nanog* and *Rex1* in iPSCs. A GSEA analysis identifies several sets of genes belonging to important biological pathways that are significantly upregulated (97 sets of genes) or downregulated (39 sets of genes) in iPSCs compared to NSCs confirming that the acquisition of a pluripotent state implies global transcriptome changes, including changes in the expression of imprinted genes.

Imprinted genes are a group of genes expressed monoallellicaly from either the maternally or the paternally inherited chromosome (Tucci et al., 2019). Although genomic imprinting is relatively stable in somatic cells, it has been described that iPSCs tend to lose imprinting similar to mouse ESCs and aberrant silencing of imprinted genes during iPSC reprogramming leads to altered tissue development (Yagi et al., 2019). Consistently, our RNAseq data in iPSCs and NSCs showed that around 50% of all imprinted genes analysed altered their transcriptional state, suggesting that regulation of imprinted gene expression is important to NSCs behaviour and reprogramming.

Several studies have shown that changes in DNA methylation patterns are essential for successful reprogramming, exemplified by the necessity for loss of promoter methylation in pluripotency genes (Takahashi and Yamanaka, 2006). In fact, methylation of CpG dinucleotides plays an important role in regulating gene transcription (Lee et al., 2014) thus if loss of DNA methylation is not achieved, cells will be only partially reprogrammed (Mikkelsen et al., 2008). Moreover, it has been described that the extent of DNA methylation remodelling is dependent on the choice of reprogramming factors used (Planello et al., 2014). We demonstrate here that the remodelling of the epigenetic profile during the reprogramming of NSCs into iPSCs with two factors is critical for pluripotency induction, as important changes in the methylation landscape of NSCs need to happen to obtain true pluripotent cells. Global DNA methylation changes were observed using MeDIP sequencing. Importantly, methylome analysis in iPSCs showed global loss of methylation in the vast majority of DMRs analysed, correlating with the global induction of gene expression in iPSCs observed by RNAseq. Our work highlights the important role of DNA demethylation in the acquisition of a pluripotent state. *Nanog* expression has been related to the inhibition of global DNA methylation (Theunissen et al., 2011) suggesting its role in methylome re-establishment. When we analysed in detail the methylation profile in pluripotency-associated genes, we also observed hypomethylation, especially at the promoter levels, whereas an increase of methylation is exhibited in neural-associated genes in accordance with their repression.

It is well described that DNA methylation is essential for establishment and maintenance of imprinting (Bartolomei and Ferguson-Smith, 2011). Indeed, germline DMRs are essential to establish monoallelic expression and secondary somatic DMRs play important roles in imprinting maintenance (Ferguson-Smith, 2011). More than 16 imprinted clusters are associated with maternal-specific methylation. For these loci, DNA methylation is found at promoters of protein-coding genes or non-coding RNA genes (Bartolomei and Ferguson-Smith, 2011). In contrast, paternally methylated ICRs (there are only 4 described) are located in intergenic regions (Bartolomei and Ferguson-Smith, 2011). Naïve iPSCs share several features with early mammalian embryos, such as global genome hypomethylation, which is also accompanied by a widespread loss of DNA methylation at imprinted loci (Perrera and Martello, 2019). Consistently, our findings reveal that the majority of the DMRs at the imprinted clusters are hypomethylated. Indeed, more than 80% of the maternally methylated DMRs show hypomethylation, whereas only 30% of paternally methylated DMRs are affected. These epigenetic changes at the maternally methylated DMRs correlate with changes in the expression levels of several imprinted genes within these clusters, whereas no significant changes were observed in the expression of genes at the paternally methylated clusters. Consequently, our results suggest that, genomic imprinting is finely regulated during development to acquire a specific somatic imprinting pattern in adult NSCs, essential for multipotency maintenance. Methylation changes at the DMRs within the imprinting clusters suggest that imprinting status of these clusters is altered in iPSCs.

Extended culture exposes cells to the risk of accumulation of genetic defects, allowing the loss of the epigenetic memory of the cell of origin (Nishino et al., 2011) as exemplified by the dynamics of X chromosome inactivation in female human iPSCs (Tchieu et al., 2010). Given that some imprinted genes are regulators of growth, it has been proposed that the loss of the imprinting status in iPSCs might also occur during their expansion in vitro, because of the advantage conferred to the cells (Perrera and Martello, 2019). This loss of DNA methylation results in some cases in the biallelic expression of the genes controlled by the DMR (Giulitti et al., 2019). Thuş loss of imprinting has been often measured in terms of loss of DNA methylation, however this does not always result in the loss of monoallelic expression, because other mechanisms (e.g, histone repressive marks) besides DNA methylation can regulate the expression of the silenced allele (Inoue et al., 2017). Moreover, it has been described that DNA methylation, when severely lost at an imprinted DMR is not restored after differentiation of iPSCs (Bar et al., 2017).

In conclusion, loss of imprinting is observed in somatic cells in culture but more frequently in iPSCs. Our study reveals that this is also the case in iPSCs generated from adult NSCs, indicating that the reprogramming process contributes to the loss of imprinting during the acquisition of a pluripotent state. Many questions remain regarding how mechanistically genomic imprinting is established, maintained and then erased during reprogramming. Therefore, iPSCs represent an ideal tool for the study of the regulation of genomic imprinting and, at the same time, a better understanding of the mechanisms controlling imprinted gene expression in iPSCs will improve the safety of their potential clinical applications.

## Methods

### Animals and *in vivo* manipulations

The experiments were conducted using C57BL/6 wild-type mice. Specifically, homozygous immunosuppressed *Nude* (*NU/J)* mice obtained from Jackson were used for the teratoma formation assays. All experiments involving animals were conducted in compliance with ethical regulations for animal testing and research. Moreover, these experiments were carried out following protocols approved by the ethics committee of the University of Valencia (Spain).

### Neurosphere cultures and reprogramming of NSCs into iPSCs

Adult 2-to 4-months-old mice were killed by cervical dislocation. To initiate each independent culture, the brains of two different animals were dissected and the regions containing the SVZ were isolated from each hemisphere and washed in Earle’s balanced salt solution (EBSS; Gibco). Tissues were transferred to EBSS containing 1.0 mg ml^-1^ papain (Worthington DBA), 0.2 mg ml^-1^ L-cystein (Sigma), 0.2 mg ml^-1^ EDTA (Sigma) and incubated for 20 min at 37°C. Tissue was then rinsed in EBSS, transferred to Dulbecco’s modified Eagle’s medium (DMEM)/F12 medium (1:1 v/v; Life Technologies) and carefully triturated with a fire-polished Pasteur pipette to a single cell suspension. Isolated cells were collected by centrifugation, resuspended in DMEM/F12 medium containing 2 mM L-glutamine (Gibco), 0.6% glucose (Panreac), 9.6 g ml^-1^ putrescine (Sigma), 6.3 ng ml^-1^ progesterone (Sigma), 5.2 ng ml^-1^ sodium selenite (Sigma), 0.025 mg ml^-1^ insulin (Sigma), 0.1 mg ml^-1^ transferrin (Sigma), 2 µg ml^-1^ heparin (sodium salt, grade II; Sigma) and supplemented with 20 ng ml^-1^ epidermal growth factor (EGF; Invitrogen) and 10 ng ml^-1^ fibroblast growth factor (FGF; Sigma) (Belenguer et al., 2016). Neurospheres were allowed to develop for 6 days in a 95% air-5% CO_2_ humidified atmosphere at 37 °C. For culture expansion, cells were plated at a relatively high density (75 cell/μl) and maintained for several passages.

To generate iPSCs from adult NSCs exogenous *Oct4* together with *Klf4* (2F) was used for reprogramming as previously described (Kim et al., 2008; Kim et al., 2009). To produce retroviruses expressing *Oct4* and *Klf4*, Platinum-E (Plat-E) retroviral packing cells (Cell Biolabs) were transfected with a plasmid solution containing 1 ml of Opti-MEM^TM^ (Gibco), 60 µl of 1mg ml^-1^ polyethylenimine (PEI, Polysciences) and 20 µg of the retroviral vectors pMXs-Oct4 (#13366, Addgene), pMXs-*Klf4* (#13370, Addgene) and pMXs-Cherry (pMX-2A-CH, designed and kindly provided by Dr. Jose Manuel Torres). After 24 hours, Plat-E culture medium (high glucose DMEM containing 10% foetal bovine serum FBS, 2 mM L-glutamine, 1 μg ml^-1^ Puromycin and 10 μg ml^-1^ Blasticidin) was replaced by NSCs complete medium. Transfection efficiency was checked by Cherry expression in Plat-E cells (**Fig. S1a**). The day after retrovirus-containing supernatants were collected and filtered with a 0.45 µm nitrocellulose filter. Neurospheres grown for two days were transduced with a mixture of virus-containing supernatant (SN) as follows (volume *per* plate): 3 ml of *Oct4* SN, 3 ml of *Klf4* SN, 1 ml of *Cherry* SN and 3 ml of fresh NSCs complete medium. A control of infection was made with a mixture containing 7 ml of *Cherry* retrovirus containing medium and 3 ml of fresh complete medium. In order to enhance the efficiency of retroviral infection, retrovirus mixture was supplemented with 4 µg ml^-1^ of Polybrene (Sigma). NSCs were then incubated for 14-18 hours at 37°C in a humidified incubator. Infected NSC medium was then replaced with fresh complete medium and neurospheres were allowed to develop for 6 days (**Fig. 1b**). The mouse fibroblast cell line SNL (Cell Biolabs) was used as feeder cells during the reprogramming process of infected NSCs. SNL feeder cells were first mitotically inactivated by treatment with 4 µg ml^-1^ of Mitomycin C (Sigma) for 2-4 hours. Plates were treated with 0.1% of gelatin (Sigma) at 37°C for at least 20 min and then mitomyzed SNLs were plated at high density (2.5×10^6^ cell/plate) in gelatine-treated plates (day 7). Six days after transduction, neurospheres were dissociated with Accutase® and 1,5×10^5^ of infected NSCs were re-plated on SNL *feeder* cells with reprogramming medium (RM): Glasgow Minimum Essential Medium (GMEM) containing 15% FBS, 2 mM L-glutamine, 1 mM Sodium pyruvate (Gibco) and 1 μM Leukaemia Inhibitory Factor (LIF). RM medium was changed every other day until Stage-Specific Embryonic Antigen-1 (SSEA-1; also known as CD15) positive colonies appeared (pre-iPSCs), checked by staining with StainAlive SSEA-1 Antibody (DyLight 488) (Stemgent®, 1:100 dilution) (**Fig. 1g** and **Fig. S1a**). Reprogramming medium was replaced with 2i/LIF medium (Neurobasal containing B27 supplement; Gibco, 2 mM L-glutamine, 1 mM Sodium pyruvate, 1 mg ml^-1^ Transferrin, 50 μM Insulin, 16 μg ml^-1^ Putrescine, 60 ng ml^-1^ Progesterone, 0.3 μM Sodium selenite, 50 μg ml^-1^ Bovine serum albumin, 1 μM LIF, 1 μM Imef; Millipore, and 3 μM iGSK3; Millipore) which is based on dual inhibition (2i) of mitogen-activated protein kinase (MAPK) signalling and glycogen synthase kinase-3 (GSK3) combined with LIF (Silva et al., 2008). 2i/LIF medium was changed every two days until well-defined iPSCs colonies appeared (**Fig. 1b**). To establish and expand clonal lines of iPSCs, individual colonies were isolated and plated on gelatine treated plates with 2i/LIF medium. The Embryonic Stem Cell (ESC) line E14Tg2a was used as a pluripotency positive control in the different experiments. ESCs were cultured on gelatine-treated plates and two days after plating, cells were treated with Trypsin/EDTA and re-plated following a dilution of 1:5 in RM.

### Embryoid bodies assays

Embryoid bodies were obtained using the “*hanging drops*” method. iPSCs were treated with Accumax® (Millipore) and resuspended in EB differentiation medium (GMEM containing 10% FBS, 2 mM L-glutamine and 1 mM Sodium pyruvate). Several rows of 20 µl drops of a cell suspension at 30 cells/ul were plated using a multichannel pipet (**Fig. 2a**). Plates were incubated upside-down for 3 days at 37°C in a 5% CO_2_ humidified incubator. Plates were then inverted and EB medium was added. To avoid EB attachment plates were previously treated with 0.4% poly (2-HEMA) solution (Sigma) prepared in Ethanol:Acetone (1:1). EBs were incubated for 4 more days and then plated on gelatine-treated plates for 3 more days before analysis (**Fig. 2a**).

### Teratoma formation and analysis

To evaluate the capacity of iPSCs to generate teratomas, mouse iPSCs cultures were collected by treatment with Accumax®. iPSCs were washed in PBS and resuspended in PBS supplemented with 30% Matrigel® (Corning®) (Prokhorova et al., 2009). Cells were kept on ice and drawn into a 1-mL syringe immediately before injection. Approximately 1.5×10^6^ cells/200 μL of solution were injected in the dorso-lateral area of the subcutaneous space on both sides of the mice back. Teratomas were allowed to develop for 15-20 days when the size of the teratomas was approximately 1.5-2 cm. Mice were sacrificed by cervical dislocation and teratomas were extracted for analysis. For teratoma analysis, samples were fixed in 4% paraformaldehyde (PFA) overnight at 4°C with shaking. Samples were embedded in paraffin and teratoma samples were serially sectioned into 7 µm sections using a microtome (Leica). Slices were stained with haematoxylin and eosin and cell types from the three embryonic layers were identified under the optic microscope (Nikon Eclipse Ni).

### Karyotype of iPSCs

To perform the karyotype analysis, cell division was inhibited using 0.6 µg ml^-1^ of KarioMAX® Colcemid (Gibco) at 37°C. After 2 hours, culture medium was removed and 0.85% sodium citrate, previously warmed at 37°C, was added. A cell Scraper (Biofil®) was used to raise the cells. Cell suspension was transferred to a 15 ml conical tube and incubated at 37°C for 15 min. After that, 10 drops of cold Carnoy fixative (Methanol-Acetic acid, 3:1; company?) were added to the suspension and softly mixed using a Pasteur pipette. Samples were washed several times with 5 ml of cold Carnoy solution and, after centrifugation (10 min 300 g), pellets were resuspended in 2 drops of Carnoy fixative. Cells extensions were made in microscope slides followed by heat fixation. Samples were stained with Leishmańs stain (Sigma). The number of chromosomes was determined under the optic microscope (Nikon Eclipse Ni).

### Immunocytochemistry and alkaline phosphatase (AP) staining

NSCs, iPSCs and EBs were fixed for staining with 4% PFA 0.1M PBS for 15 min and immunocytochemistry performed as previously described (Belenguer et al., 2016). Primary and secondary antibodies and dilutions used are listed in **Table S1** and **Table S2** respectively. DAPI (1µg ml^-1^) was used to counterstain DNA. Samples were photographed and analysed using an FV10i confocal microscope (Olympus). Alkaline phosphatase detection method was used in reprogrammed cells to check the presence of iPSCs after one month in 2i/LIF medium on *feeders*. Cells were fixed with cold methanol for 2 min and washed three times with 0.1M Tris-HCl pH 8.5 buffer. Samples were incubated with the “*staining solution*” which contained 0.1 mg ml^-1^ Naphtol phosphate (Sigma), 0.5% Dimethylformamide (Sigma) and 0.6 mg ml^-1^ Fast Red Salt (Sigma) in 0.1M Tris-HCl pH 8.5. When red precipitate appeared, cells were washed with 0.1M Tris-HCl and distilled water. Finally, the different plates were photographed using a dissection microscope.

### Expression studies

RNAs were extracted with RNAeasy mini kit (Qiagen) including DNase treatment, following the manufacturer’s guidelines. For qPCR, 1 μg of total RNA was reverse transcribed using random primers and RevertAid H Minus First Strand cDNA Synthesis kit (Thermo Scientific), following standard procedures. Thermocycling was performed in a final volume of 10 μl, containing 1 μl of cDNA sample and the reverse transcribed RNA was amplified by PCR with appropriate Taqman probes (**Table S3**). RT-PCR was used to measure gene expression levels normalized to *Gapdh*, the expression of which did not differ between the groups. qPCR reactions were performed in a Step One Plus cycler with Taqman Fast Advanced Master Mix (Applied Biosystems). SYBR green thermocycling was also performed in a final volume of 12 μl, containing 1 μl of cDNA sample, 0.2µM of each primer (**Table S4**) and SYBR® Premix ExTaq^TM^ (Takara) according to the manufacture instructions. A standard curve made up of doubling dilutions of pooled cDNA from the samples being assessed was run on each plate, and quantification was performed relative to the standard curve.

### RNA-seq

Library preparation and high-throughput sequencing were performed by the Central Service for Experimental Research (SCSIE) at the University of Valencia. RNA-seq libraries were generated from triplicated samples per condition using the Illumina TruSeq stranded mRNA Sample Preparation Kit v2 (company?) following the manufacturer’s protocol. The RNA-seq libraries were sequenced using Illumina NextSeq 500. Analysis of the RNA-seq data was performed by EpiDisease S.L. The sequences were aligned to the mouse genome reference (GRCm38.p5) taken from Ensembl using the Subread package in Linux. The expression of every gene per sample was measured using a gtf annotation file (GRCm38.87) taken from Ensembl and the R package Rsubread. Data that support the findings of this study have been deposited in Gene Expression Omnibus (GEO) with the accession number XXXXXX

### High-Throughput Sequencing of Immunoprecipitated Methylated DNA (MeDIP-seq)

The DNeasy Blood and Tissue Kit (Qiagen) was used to extract DNA from NSCs and iPSCs. Extraction was performed following the manufacturer’s instructions. Briefly, a reducing agent (DTT) and Proteinase K were added to the lysate buffer; samples were incubated overnight at 55°C. Samples were eluted in 100 μl and the concentration of DNA was measured using a Nanodrop 1000. MeDIP-seq protocol was modified from Taiwo et. al. (Taiwo et al., 2012). For immunoprecipitation, 3 µg of DNA were sonicated to obtain 150-200 bp fragments and the efficiency was checked by the Bioanalyzer (Agilent). DNA libraries were prepared using NEBNext® reactives (New England Biolabs). For MeDIP, 1.5 µg of DNA was diluted in TE buffer (10 mM Tris-HCl, 1 mM EDTA, pH 7.5) and denatured for 10 min at 99°C. After that, 20 µl of 10x IP buffer (100 mM Na-Phosphate pH 7.0, 0.5% TritonX-100) and 100 µl of 5% skimmed milk buffer in 2M NaCl was added. 2 µg of 5mC antibody (Diagenode) were added to the sample and incubated for 2 h at 4°C with rotation. In parallel, 11 µl per sample of Dynabeads® M-280 sheep anti-mouse IgG (Thermo Fisher) were collected with a magnetic rack and pre-washed with 500 µl PBS-BSA (1 mg/ml BSA in 0.1 M PBS) for 2 hours at 4°C with rotation. After incubations, beads were collected with a magnet, resuspended in the original volume with 1x IP buffer (10 mM Na-Phosphate pH 7.0, 0.05% TritonX-100, 1 M NaCl) and added to the DNA samples. DNAs were incubated overnight at 4°C with rotation. The day after, beads were collected using the magnet and the supernatant (unbound fraction) was transferred to a new tube. Beads were washed three times with 500 µl of 1x IP buffer for 10 min with rotation. After the final wash, bound and unbound fractions were treated with 0.3 mg ml^-1^ of Proteinase K (Roche) in digestion buffer (50 mM Tris-HCl pH 8.0, 10 mM EDTA, 0.5% SDS) and incubated at 55°C for 30 min on a shaking heating block. Samples were purified using MiniElute PCR purification kit (Qiagen) and eluted in 10 µl of elution buffer. In order to calculate 5mC enrichment in the bound fraction, quantitative PCRs for unmethylated and methylated regions were done from bound and unbound fractions. Primers used to evaluate MeDIP efficiency were for methylated regions, Meth-F: 5’-CATGGCCCACAAAGTAATAAAA-3’ and Meth-R: 5’-AACGACTTACAACGAGCTCAAA-3. Primers for unmethylated regions were, Unmeth-F: 5’-GGCTAGAACTGACCAGACAGAC-3’ and Unmeth-R: 5’-ATCTGTAGCCAATCCTAGAGCA-3’. Enrichment should be of at least 25x, specificity should be more than 95% and unmethylated recovery should be less than 1%. The high-throughput sequencing of the samples was done using HiSeq2000 (Illumina, Inc) technology. For analysis of MeDIP-seq data, adaptor sequences were trimmed and reads were filtered using afterQC (Chen et al., 2017). Reads were removed if: one of the read pair had a polyX run greater than 35bp long; if the phred score was less than 20; if 5 bases or more were called as ‘N’ or if the trimmed read was shorter than 35bp. Reads were then mapped to mm10 using BWA-MEM. Unmapped and unpaired reads were removed using Samtools. MeDIP data was analysed in R using the MEDIPS package (Lienhard et al., 2014). A sliding window of 250 bp was used. A minimum read depth of 20 across all samples; CpG density normalisation; correction for read depth and Benjamini Hochberg multiple testing correction was applied. There were no regions of significantly altered methylation after correction for multiple testing. PCA analysis was conducted in python using scikit-learn and plotted using Matplotlib.

### DNA methylation analysis and pyrosequencing

DNA methylation level was quantified using bisulfite conversion and pyrosequencing. The DNA was bisulfite-converted using EZ DNA Methylation-Gold^TM^ kit (Zymo research) in accordance with the manufacturés protocol. Specifically, for the different DMRs, bisulfite-converted DNA was amplified by PCR with specific primer pairs (**Table S4**). PCRs were carried out in 20 µl, with 2U HotStar Taq polymerase (Qiagen), PCR Buffer 10x (Qiagen), 0.2 mM dNTPs and 400 mM primers. PCR conditions were: 96°C for 5 min, followed by 39 cycles of 94°C for 30 s, 54°C for 30 s and 72°C for 1 min. For pyrosequencing analysis, a biotin-labelled primer was used to purify the final PCR product using sepharose beads. The PCR product was bound to Streptavidin Sepharose High Performance (GE Healthcare), purified, washed with 70% ethanol, denatured with 0.2 N NaOH and washed again with 10 mM Tris-acetate. Pyrosequencing primer (400 mM) was then annealed to the purified single-stranded PCR product and pyrosequencing was performed using the PyroMark Q96MD pyrosequencing system using PyroMark® reactives (Qiagen).

### Statistical analysis

All statistical tests were performed using the GraphPad Prism Software, version 7.00 for Windows. Data were first tested for normality with a Shapiro-Wilk test. The significance of the differences between groups were evaluated with adequate statistical tests for each comparison. For data that passed normality tests: when analysing only one variable, a paired t-test was used for comparing two groups and one-way ANOVA followed by Tukey’s post-hoc test for three or more groups. When two variables were analyse, two-way ANOVA followed by Tukey post-hoc test was used. For data groups that did not pass normality, Wilcoxon or Mann–Whitney nonparametric tests were performed, depending on whether samples were paired or not, respectively. For variables with more than two categories, Kruskal– Wallis or Friedman tests (for unpaired or paired data, respectively) were used followed by a Benjamini, Krieger and Yekutieli post-hoc test. When comparisons were performed with relative values (percentages), data were previously normalized by using arcsin root transformation. Values of P<0.05 were considered statistically significant. Data are presented as the mean ± standard error of the mean (s.e.m.) and the number of experiments performed with independent cultures or animals (n) and P-values are indicated in the figures. Box and whisker plots show the mean (+), median (horizontal line in box) and maximum and minimum values (whiskers).

## Acknowledgements

We firstly would like to thank Dr. Isabel Fariñas, Dr. Anne Ferguson-Smith and Dr. Ángel Raya and their groups for technical support and discussion of the data. This work was supported by grants from Ministerio de Ciencia e Innovación (PID2019-110045GB-I00, PID2022-142734OB-I00 and EUR2023-143479), Generalitat Valenciana (AICO/2020/367) and Fundación BBVA to SRF. RML (CÓDIGO) and LLC (PRE2020-094137) were funded by the Spanish Formación de Personal Investigador (FPI) fellowship program. EJV was funded by the Spanish Formación de Profesorado Universitario fellowship program (FPU20/00795). ALU was funded by the Generalitat Valenciana fellowship program (ACIF/2016/381). Open Access funding provided by the Ministerio de Ciencia e Innovación.

## Author Contribution

RML, ALU, LLC, EJV and ALP performed most of the experiments. LLC helped with the gene expression analysis. ALP helped to develop the reprogramming protocol. MI helped with methylation studies. JP performed the bioinformatics analysis-ER performed the analysis of the MeDIP-seq data. SRF initiated, designed and led the study, and wrote the manuscript. All authors contributed to experimental design, data analysis, discussion and writing of the paper.

## Data availability

All relevant data can be found within the article and its supplementary information.

---

**Figure S1. The reprogramming of adult NSCs generates iPSCs that express pluripotency genes. (a)** Phase contrast and fluorescent images for Cherry (red) of Plat-E, post-infected NSCs (NSCs-PI) and pre-iPSCs. An immunofluorescent image for SSEA-1 (green) in pre-iPSCs cells is also shown. **(b)** Immunocytochemistry images for OCT4 (green), OLIG2 (blue) and Cherry (red) in iPSCs. **(c)** qPCR of the pluripotency-related gene *Zfp42* in adult NSCs, pre-iPSCs and iPSCs. ESCs were used as a positive control. **(d)** Alkaline phosphatase (AP) staining in iPSCs generated from NSCs. DAPI was used to counterstain nuclei. *Gapdh* was used to normalize gene expression data. Scale bars in a: 100 µm; high magnification images in a and d: 10 µm; in b: 20 µm. All error bars show s.e.m. P-values and number of samples are indicated.

**Figure S2. EBs generated from ESCs express markers from the three germinal layers. (a)** qPCR for *Kdr1* (mesoderm), *Foxa2* (endoderm) and *Cer1* (ectoderm) genes in ESCs and in embryoid bodies (EBs) derived from ESCs. **(b)** Phase contrast image of EBs derived from ESCs (left panel). Immunocytochemistry images for α-fetoprotein (green) and for α-SMA in EBs differentiated from the ESCs. DAPI was used to counterstain nuclei. *Gapdh* was used to normalize gene expression data. Scale bars in b: 50 µm in phase contrast images and 10 µm in fluorescence images. All error bars show s.e.m. P-values and number of samples are indicated.

**Figure S3. Reprogramming of NSCs causes global changes in gene expression. (a)** Pie graphs showing the percentage of genes that are upregulated (light blue) or dowregulated (dark blue) in iPSCs compared to NSCs. The percentage of imprinted genes unaltered are also shown. **(b)** Gene set enrichment analysis (GSEA) representing in dark blue the number of downregulated genes in pathways with more than 35 genes differentially expressed. GSEA representing in light blue the number of upregulated genes in pathways with more than 50 differentially expressed genes. **(c)** Gene ontology (GO) analysis representing the fold enrichment of genes between iPSCs and NSCs classified in different categories according to biological processes and cellular components.

**Figure S4. Methylation profiles at the promoters of pluripotency and neural genes are altered in iPSCs. (a)** qPCR for the maternally expressed genes *Cdkn1c*, *Gnasxl* and *Igf2r* (pink) and for the paternally expressed gene *Peg3, Kcnqlot1l* and *Mcts2* (blue) in NSCs and iPSCs. **(b)** DNA methylation profile analysis of *Nanog, Oct4* and *Zfp42* (left panels) and *Nestin*, *Zic1* and *Olig2* (right panels) at the promoter regions (grey) by MeDIP-seq in NSCs and iPSCs. A schematic representation of each gene is included. *Gapdh* was used to normalize expression data. All error bars show s.e.m. of at least 4 samples. n.s.: no significant.

**Figure S5. Changes in the expression of imprinted genes in iPSCs correlate with hypomethylation at the maternally methylated DMRs (a)** Schematic of the maternally methylated clusters *Kcnq1ot1* DMR, *Snrpn* DMR and *Grb10* DMR. Expression of genes within these clusters and methylation profiles of the DMRs are shown. Downregulated genes are indicated in dark blue and upregulated are in light blue. **(b)** Schematic of the paternally methylated imprinting clusters showing the IG-DMR and the H19 ICR. Log_2_ fold change of the expression of imprinted genes within these clusters is shown.

## References

Aasen, T., Raya, A., Barrero, M. J., Garreta, E., Consiglio, A., Gonzalez, F., Vassena, R., Bilić, J., Pekarik, V., Tiscornia, G., et al. (2008). Efficient and rapid generation of induced pluripotent stem cells from human keratinocytes. Nat. Biotechnol. 26, 1276–1284.

Aoi, T., Yae, K., Nakagawa, M., Ichisaka, T., Okita, K., Takahashi, K., Chiba, T. and Yamanaka, S. (2008). Generation of Pluripotent Stem Cells from Adult Mouse Liver and Stomach Cells. Science 321, 699–702.

Apostolou, E., Ferrari, F., Walsh, R. M., Bar-Nur, O., Stadtfeld, M., Cheloufi, S., Stuart, H. T., Polo, J. M., Ohsumi, T. K., Borowsky, M. L., et al. (2013). Genome-wide Chromatin Interactions of the Nanog Locus in Pluripotency, Differentiation, and Reprogramming. Cell Stem Cell 12, 699–712.

Arez, M., Eckersley-Maslin, M., Klobučar, T., Lopes, J. von G., Krueger, F., Mupo, A., Raposo, A. C., Oxley, D., Mancino, S., Gendrel, A.-V., et al. (2022). Imprinting fidelity in mouse iPSCs depends on sex of donor cell and medium formulation. Nat. Commun. 13, 5432.

Bar, S., Schachter, M., Eldar-Geva, T. and Benvenisty, N. (2017). Large-Scale Analysis of Loss of Imprinting in Human Pluripotent Stem Cells. Cell Rep. 19, 957–968.

Barlow, D. P. and Bartolomei, M. S. (2014). Genomic imprinting in mammals. Cold Spring Harb. Perspect. Biol. 6, a018382–a018382.

Bartolomei, M. S. and Ferguson-Smith, A. C. (2011). Mammalian Genomic Imprinting. Cold Spring Harb. Perspect. Biol. 3, a002592.

Belenguer, G., Domingo-Muelas, A., Ferrón, S. R., Morante-Redolat, J. M. and Fariñas, I. (2016). Isolation, culture and analysis of adult subependymal neural stem cells. Differentiation 91, 28–41.

Cantone, I. and Fisher, A. G. (2017). Human X chromosome inactivation and reactivation: implications for cell reprogramming and disease. Philos. Trans. R. Soc. Lond. Ser. B, Biol. Sci. 372, 20160358.

Carter, A. C., Davis-Dusenbery, B. N., Koszka, K., Ichida, J. K. and Eggan, K. (2014). Nanog-Independent Reprogramming to iPSCs with Canonical Factors. Stem Cell Rep. 2, 119–126.

Chen, S., Huang, T., Zhou, Y., Han, Y., Xu, M. and Gu, J. (2017). AfterQC: automatic filtering, trimming, error removing and quality control for fastq data. BMC Bioinform. 18, 80.

Edwards, C. A. and Ferguson-Smith, A. C. (2007). Mechanisms regulating imprinted genes in clusters. Curr. Opin. Cell Biol. 19, 281–289.

Eminli, S., Utikal, J., Arnold, K., Jaenisch, R. and Hochedlinger, K. (2008). Reprogramming of Neural Progenitor Cells into Induced Pluripotent Stem Cells in the Absence of Exogenous Sox2 Expression. STEM CELLS 26, 2467–2474.

Ferguson-Smith, A. C. (2011). Genomic imprinting: the emergence of an epigenetic paradigm. Nat. Rev. Genet. 12, 565–575.

Ferrón, S. R., Charalambous, M., Radford, E., McEwen, K., Wildner, H., Hind, E., Morante-Redolat, J. M., Laborda, J., Guillemot, F., Bauer, S. R., et al. (2011). Postnatal loss of Dlk1 imprinting in stem cells and niche-astrocytes regulates neurogenesis. Nature 475, 381–385.

Ferrón, S. R., Radford, E. J., Domingo-Muelas, A., Kleine, I., Ramme, A., Gray, D., Sandovici, I., Constancia, M., Ward, A., Menheniott, T. R., et al. (2015). Differential genomic imprinting regulates paracrine and autocrine roles of IGF2 in mouse adult neurogenesis. Nat. Commun. 6, 8265.

Giulitti, S., Pellegrini, M., Zorzan, I., Martini, P., Gagliano, O., Mutarelli, M., Ziller, M. J., Cacchiarelli, D., Romualdi, C., Elvassore, N., et al. (2019). Direct generation of human naive induced pluripotent stem cells from somatic cells in microfluidics. Nat. Cell Biol. 21, 275–286.

Hanna, J., Carey, B. W. and Jaenisch, R. (2008). Reprogramming of Somatic Cell Identity. Cold Spring Harb. Symp. Quant. Biol. 73, 147–155.

Hochedlinger, K. and Jaenisch, R. (2015). Induced Pluripotency and Epigenetic Reprogramming. Cold Spring Harb. Perspect. Biol. 7, a019448.

Höpfl, G., Gassmann, M. and Desbaillets, I. (2004). Differentiating embryonic stem cells into embryoid bodies. Methods Mol. Biol. (Clifton, NJ) 254, 79–98.

Hotta, A. and Ellis, J. (2008). Retroviral vector silencing during iPS cell induction: An epigenetic beacon that signals distinct pluripotent states. J. Cell. Biochem. 105, 940–948.

Inoue, A., Jiang, L., Lu, F., Suzuki, T. and Zhang, Y. (2017). Maternal H3K27me3 controls DNA methylation-independent imprinting. Nature 547, 419–424.

Janiszewski, A., Talon, I., Chappell, J., Collombet, S., Song, J., Geest, N. D., To, S. K., Bervoets, G., Marin-Bejar, O., Provenzano, C., et al. (2019). Dynamic reversal of random X-Chromosome inactivation during iPSC reprogramming. Genome Res. 29, 1659–1672.

Kim, J. B., Zaehres, H., Wu, G., Gentile, L., Ko, K., Sebastiano, V., Araúzo-Bravo, M. J., Ruau, D., Han, D. W., Zenke, M., et al. (2008). Pluripotent stem cells induced from adult neural stem cells by reprogramming with two factors. Nature 454, 646–650.

Kim, J. B., Zaehres, H., Araúzo-Bravo, M. J. and Schöler, H. R. (2009). Generation of induced pluripotent stem cells from neural stem cells. Nat. Protoc. 4, 1464–1470.

Kim, M. J., Choi, H. W., Jang, H. J., Chung, H. M., Arauzo-Bravo, M. J., Schöler, H. R. and Do, J. T. (2013). Conversion of genomic imprinting by reprogramming and redifferentiation. J. Cell Sci. 126, 2516–2524.

Kuttler, F. and Mai, S. (2006). Genome and Disease. Genome Dyn. 1, 171–190.

Lassi, G. and Tucci, V. (2019). Genomic imprinting and the control of sleep in mammals. Curr. Opin. Behav. Sci. 25, 77–82.

Lee, H. J., Hore, T. A. and Reik, W. (2014). Reprogramming the Methylome: Erasing Memory and Creating Diversity. Cell Stem Cell 14, 710–719.

Lee, H. J., Choi, N. Y., Lee, S.-W., Ko, K., Hwang, T. S., Han, D. W., Lim, J., Schöler, H. R. and Ko, K. (2016). Epigenetic alteration of imprinted genes during neural differentiation of germline-derived pluripotent stem cells. Epigenetics 11, 177–183.

Lienhard, M., Grimm, C., Morkel, M., Herwig, R. and Chavez, L. (2014). MEDIPS: genome-wide differential coverage analysis of sequencing data derived from DNA enrichment experiments. Bioinformatics 30, 284–286.

Liu, L., Luo, G.-Z., Yang, W., Zhao, X., Zheng, Q., Lv, Z., Li, W., Wu, H.-J., Wang, L., Wang, X.-J., et al. (2010). Activation of the Imprinted Dlk1-Dio3 Region Correlates with Pluripotency Levels of Mouse Stem Cells. J. Biol. Chem. 285, 19483–19490.

Liu, X., Zheng, K., Zhao, X., Xu, X., Yang, A., Yi, M., Tao, H., Xie, B., Qiu, M. and Yang, J. (2020). Chromosomal aberration arises during somatic reprogramming to pluripotent stem cells.

Mikkelsen, T. S., Hanna, J., Zhang, X., Ku, M., Wernig, M., Schorderet, P., Bernstein, B. E., Jaenisch, R., Lander, E. S. and Meissner, A. (2008). Dissecting direct reprogramming through integrative genomic analysis. Nature 454, 49–55.

Montalbán-Loro, R. (2015). Epigenetic regulation of stemness maintenance in the neurogenic niches. World J. Stem Cells 7, 700.

Montalbán-Loro, R., Lassi, G., Lozano-Ureña, A., Perez-Villalba, A., Jiménez-Villalba, E., Charalambous, M., Vallortigara, G., Horner, A. E., Saksida, L. M., Bussey, T. J., et al. (2021). Dlk1 dosage regulates hippocampal neurogenesis and cognition. Proc. Natl. Acad. Sci. 118, e2015505118.

Nashun, B., Hill, P. W. and Hajkova, P. (2015). Reprogramming of cell fate: epigenetic memory and the erasure of memories past. EMBO J. 34, 1296–1308.

Nazor, K. L., Altun, G., Lynch, C., Tran, H., Harness, J. V., Slavin, I., Garitaonandia, I., Müller, F.-J., Wang, Y.-C., Boscolo, F. S., et al. (2012). Recurrent Variations in DNA Methylation in Human Pluripotent Stem Cells and Their Differentiated Derivatives. Cell Stem Cell 10, 620–634.

Nishino, K., Toyoda, M., Yamazaki-Inoue, M., Fukawatase, Y., Chikazawa, E., Sakaguchi, H., Akutsu, H. and Umezawa, A. (2011). DNA Methylation Dynamics in Human Induced Pluripotent Stem Cells over Time. PLoS Genet. 7, e1002085.

Ohhata, T. and Wutz, A. (2013). Reactivation of the inactive X chromosome in development and reprogramming. Cell. Mol. Life Sci. 70, 2443–2461.

Parry, A., Rulands, S. and Reik, W. (2021). Active turnover of DNA methylation during cell fate decisions. Nat. Rev. Genet. 22, 59–66.

Pasque, V. and Plath, K. (2015). X chromosome reactivation in reprogramming and in development. Curr. Opin. cell Biol. 37, 75–83.

Perez, J. D., Rubinstein, N. D. and Dulac, C. (2016). New Perspectives on Genomic Imprinting, an Essential and Multifaceted Mode of Epigenetic Control in the Developing and Adult Brain. Annu. Rev. Neurosci. 39, 347–84.

Perrera, V. and Martello, G. (2019). How Does Reprogramming to Pluripotency Affect Genomic Imprinting? Front. Cell Dev. Biol. 7, 76.

Planello, A. C., Ji, J., Sharma, V., Singhania, R., Mbabaali, F., Müller, F., Alfaro, J. A., Bock, C., Carvalho, D. D. D. and Batada, N. N. (2014). Aberrant DNA methylation reprogramming during induced pluripotent stem cell generation is dependent on the choice of reprogramming factors. Cell Regen. 3, 4.

Polo, J. M., Liu, S., Figueroa, M. E., Kulalert, W., Eminli, S., Tan, K. Y., Apostolou, E., Stadtfeld, M., Li, Y., Shioda, T., et al. (2010). Cell type of origin influences the molecular and functional properties of mouse induced pluripotent stem cells. Nat. Biotechnol. 28, 848–855.

Prokhorova, T. A., Harkness, L. M., Frandsen, U., Ditzel, N., Schrder, H. D., Burns, J. S. and Kassem, M. (2009). Teratoma Formation by Human Embryonic Stem Cells Is Site Dependent and Enhanced by the Presence of Matrigel. Stem Cells Dev. 18, 47–54.

Przyborski, S. A. (2005). Differentiation of human embryonic stem cells after transplantation in immune-deficient mice. Stem cells (Dayt., Ohio) 23, 1242–50.

SanMiguel, J. M. and Bartolomei, M. S. (2018). DNA methylation dynamics of genomic imprinting in mouse development†. Biol. Reprod. 99, 252–262.

Silva, J., Barrandon, O., Nichols, J., Kawaguchi, J., Theunissen, T. W. and Smith, A. (2008). Promotion of Reprogramming to Ground State Pluripotency by Signal Inhibition. PLoS Biol. 6, e253.

Smallwood, S. A. and Kelsey, G. (2012). De novo DNA methylation: a germ cell perspective. Trends Genet. 28, 33– 42.

Spelke, D. P., Ortmann, D., Khademhosseini, A., Ferreira, L. and Karp, J. M. (2010). Methods for embryoid body formation: the microwell approach. Methods Mol. Biol. (Clifton, NJ) 690, 151–62.

Stadtfeld, M., Brennand, K. and Hochedlinger, K. (2008). Reprogramming of pancreatic beta cells into induced pluripotent stem cells. Curr. Biol.: CB 18, 890–4.

Stadtfeld, M., Apostolou, E., Akutsu, H., Fukuda, A., Follett, P., Natesan, S., Kono, T., Shioda, T. and Hochedlinger, K. (2010). Aberrant silencing of imprinted genes on chromosome 12qF1 in mouse induced pluripotent stem cells. Nature 465, 175–181.

Takahashi, K. and Yamanaka, S. (2006). Induction of Pluripotent Stem Cells from Mouse Embryonic and Adult Fibroblast Cultures by Defined Factors. Cell 126, 663–676.

Takahashi, N., Gray, D., Strogantsev, R., Noon, A., Delahaye, C., Skarnes, W. C., Tate, P. H. and Ferguson-Smith, A. C. (2015). ZFP57 and the Targeted Maintenance of Postfertilization Genomic Imprints. Cold Spring Harb. Symp. Quant. Biol. 80, 177–187.

Takikawa, S., Ray, C., Wang, X., Shamis, Y., Wu, T.-Y. and Li, X. (2013). Genomic imprinting is variably lost during reprogramming of mouse iPS cells. Stem Cell Res. 11, 861–873.

Tchieu, J., Kuoy, E., Chin, M. H., Trinh, H., Patterson, M., Sherman, S. P., Aimiuwu, O., Lindgren, A., Hakimian, S., Zack, J. A., et al. (2010). Female Human iPSCs Retain an Inactive X Chromosome. Cell Stem Cell 7, 329–342.

Theunissen, T. W., Oosten, A. L. van, Castelo-Branco, G., Hall, J., Smith, A. and Silva, J. C. R. (2011). Nanog Overcomes Reprogramming Barriers and Induces Pluripotency in Minimal Conditions. Curr. Biol. 21, 65–71.

Tian, C., Wang, Y., Sun, L., Ma, K. and Zheng, J. C. (2011). Reprogrammed mouse astrocytes retain a “memory” of tissue origin and possess more tendencies for neuronal differentiation than reprogrammed mouse embryonic fibroblasts. Protein Cell 2, 128–140.

Tucci, V., Isles, A. R., Kelsey, G., Ferguson-Smith, A. C., Group, the E. I., Tucci, V., Bartolomei, M. S., Benvenisty, N., Bourc’his, D., Charalambous, M., et al. (2019). Genomic Imprinting and Physiological Processes in Mammals. Cell 176, 952–965.

Utikal, J., Maherali, N., Kulalert, W. and Hochedlinger, K. (2009). Sox2 is dispensable for the reprogramming of melanocytes and melanoma cells into induced pluripotent stem cells. J. Cell Sci. 122, 3502–3510.

Vaz, I. M., Borgonovo, T., Kasai-Brunswick, T. H., Santos, D. S. dos, Mesquita, F. C. P., Vasques, J. F., Gubert, F., Rebelatto, C. L. K., Senegaglia, A. C. and Brofman, P. R. S. (2021). Chromosomal aberrations after induced pluripotent stem cells reprogramming. Genet. Mol. Biol. 44, e20200147.

Wernig, M., Meissner, A., Cassady, J. P. and Jaenisch, R. (2008). c-Myc Is Dispensable for Direct Reprogramming of Mouse Fibroblasts. Cell Stem Cell 2, 10–12.

Yagi, M., Kabata, M., Ukai, T., Ohta, S., Tanaka, A., Shimada, Y., Sugimoto, M., Araki, K., Okita, K., Woltjen, K., et al. (2019). De Novo DNA Methylation at Imprinted Loci during Reprogramming into Naive and Primed Pluripotency. Stem Cell Rep. 12, 1113–1128.

Ying, Q.-L., Wray, J., Nichols, J., Batlle-Morera, L., Doble, B., Woodgett, J., Cohen, P. and Smith, A. (2008). The ground state of embryonic stem cell self-renewal. Nature 453, 519–523.

