## Supplementary Information for "Alterations of genomic imprinting appear during the reprogramming of adult neural stem cells"

### Figure Supplementary 1

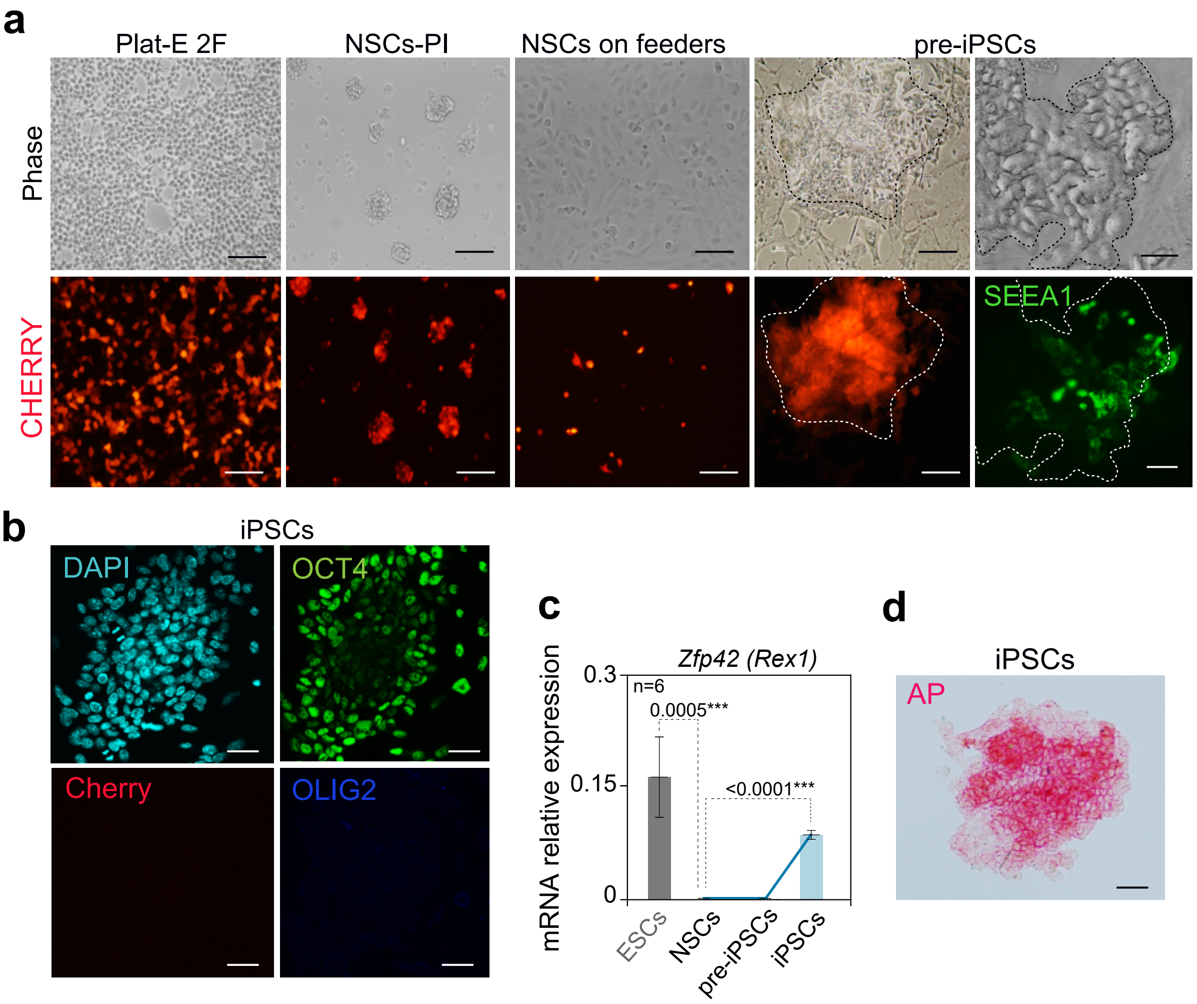

### Figure Supplementary 2

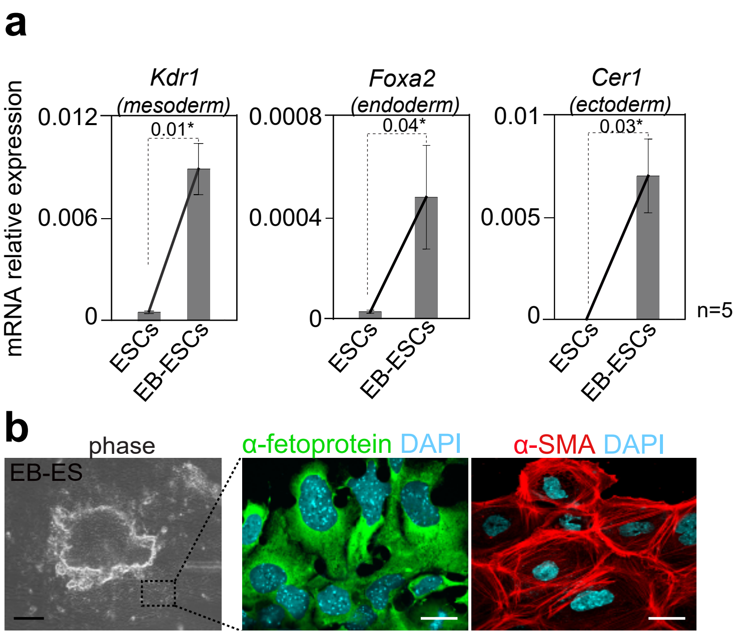

### Figure Supplementary 3

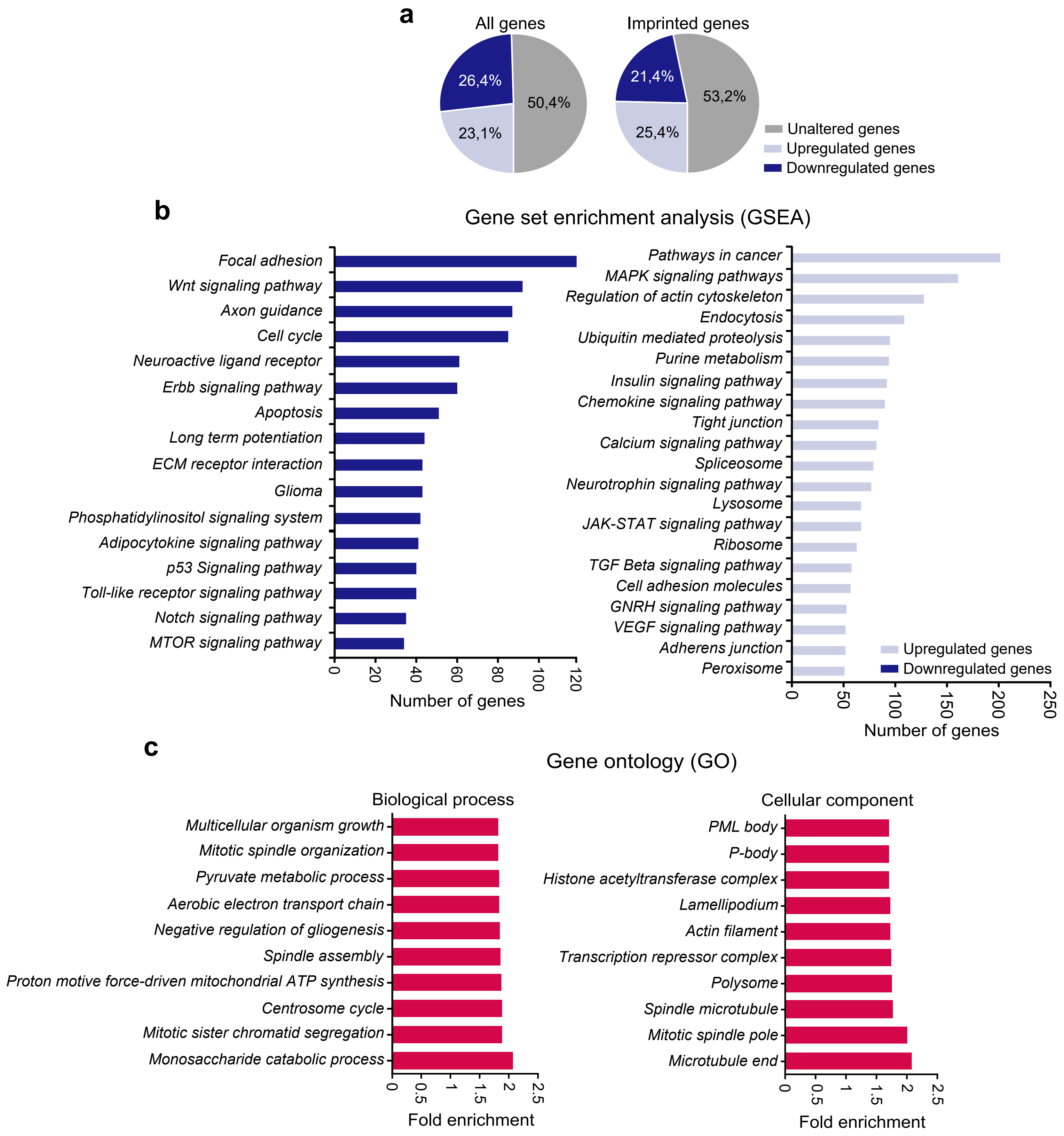

### Figure Supplementary 4

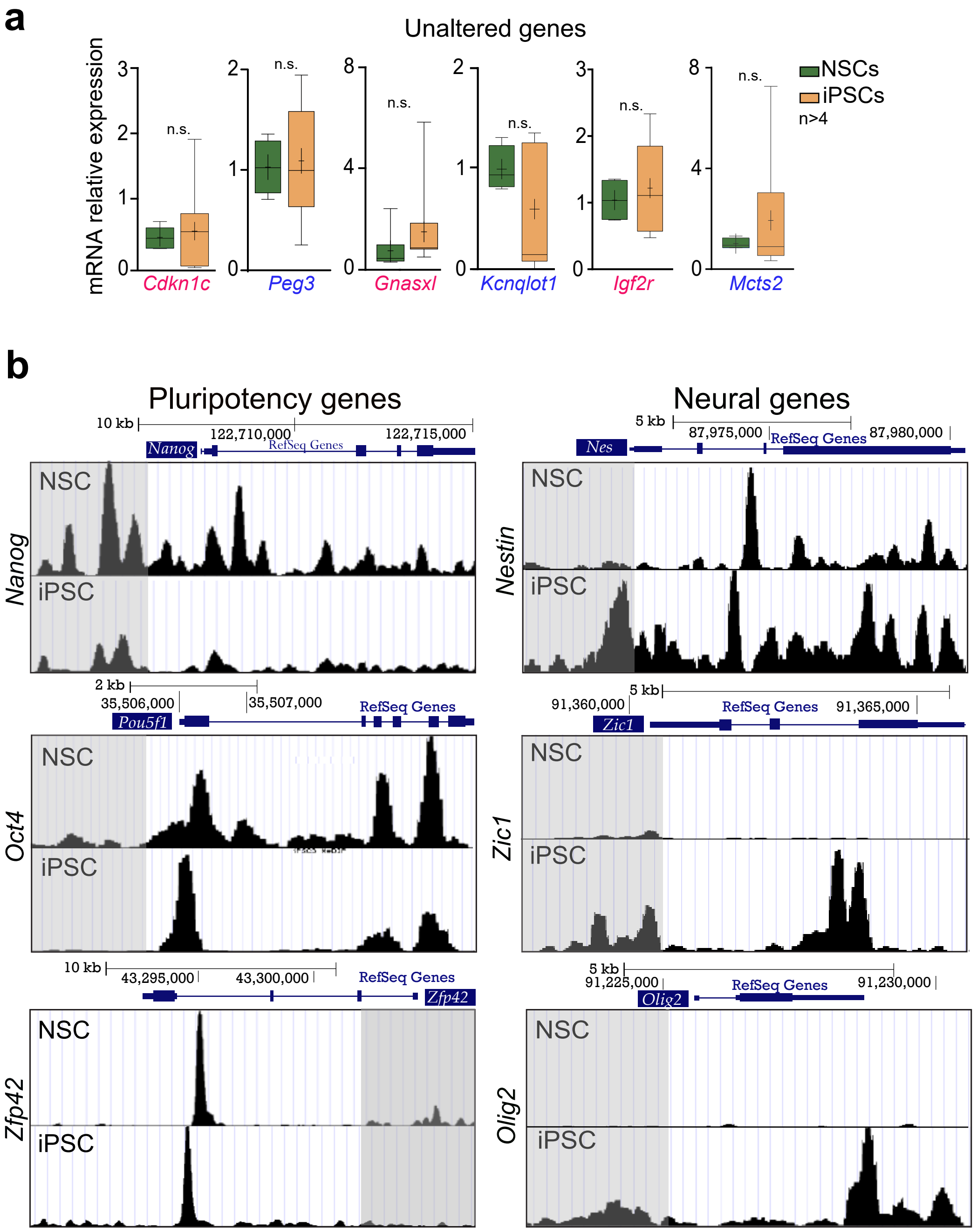

### Supplementary Tables

**Supplementary Table 1. List of primary antibodies along the different applications.**
ICC: immunocytochemistry.

| ***Primary antibody*** | ***Source*** | ***Host*** | ***Dilution*** | ***Cat. no.*** | ***Application*** |
|---|---|---|---|---|---|
| α-fetoprotein | R&D | Rabbit | 1:100 | mab1368 | ICC |
| βIII-tubulin | Covance | Mouse | 1:300 | PRB-435P | ICC |
| Brachyury | Santa Cruz | Goat | 1:500 | sc-17743 | ICC |
| GATA-4 | Santa Cruz | Goat | 1:1000 | sc-1237 | ICC |
| Nanog | Reprocell | Rabbit | 1:100 | RCAB002PF | ICC |
| OCT-4 | Santa Cruz | Rabbit | 1:200 | sc-5279 | ICC |
| OLIG2 | Millipore | Rabbit | 1:500 | AB9610 | ICC |
| α-Smooth Muscle Actin (α-SMA) | Abcam | Mouse | 1:100 | ab18147 | ICC |
| SOX2 | R&D Systems | Goat | 1:200 | AF2018 | ICC |
| SSEA1 | Santa Cruz | Mouse | 1:50 | SC-21702 | ICC |

**Supplementary Table 2. List of secondary antibodies along the different application.**
ICC: immunocytochemistry.

| ***Secondary antibody*** | ***Source*** | ***Dilution*** | ***Cat. no.*** | ***Application*** |
|---|---|---|---|---|
| Alexa Fluor® 488 Donkey Anti-Goat | Jackson ImmunoResearch | 1:1000 | 705-547-003 | ICC |
| Alexa Fluor® 488 Donkey Anti-Mouse | Molecular Probes | 1:1000 | A-21202 | ICC |
| Alexa Fluor® 488 Donkey Anti-Rabbit | Jackson ImmunoResearch | 1:1000 | 711-547-003 | ICC |
| Alexa Fluor® 647 Donkey Anti-Rabbit | Jackson ImmunoResearch | 1:1000 | 711-607-003 | ICC |
| Cy3-Donkey Anti-Rabbit | Jackson ImmunoResearch | 1:2000 | 711-165-152 | ICC |
| Cy3-Donkey Anti-Mouse | Jackson ImmunoResearch | 1:2000 | 715-165-151 | ICC |
| Cy3-Donkey Anti-goat | Jackson ImmunoResearch | 1:2000 | 705-166-147 | ICC |

**Supplementary Table 3. List of TaqMan probes used.** *TaqMan probes designed by us.

| ***Gene*** | ***Taqman code (Applied Biosystems)*** | ***Gene*** | ***Taqman code (Applied Biosystems)*** |
|---|---|---|---|
| *Afp* | Mm00431715_m1 | *Ndn* | Mm02524479_s1 |
| *Cdkn1c* | Mm01272135_g1 | *Oct4* | Mm00658129_gH |
| *Cer1* | Mm00515474_m1 | *Olig2* | Mm01210556_m1 |
| *Cntn3* | Mm00500947_m1 | *Pax6* | Mm00443081_m1 |
| *c-myc* | Mm00487803_m1 | *Peg3* | Mm01337379_m1 |
| *Dio3* | Mm00548953_s1 | *Peg10* | Mm01167724_m1 |
| *Foxa2* | Mm01976556_s1 | *Peg12* | Mm00844053_s1 |
| *Gapdh* | Mm99999915_g1 | *Phlda2* | Mm00493899_g1 |
| *Grb10* | Mm01180443_m1 | *Plagl1* | Mm00494251_m1 |
| *H19* | Mm01156721_g1 | *Ppp1r9a* | Mm00725102_m1 |
| *Igf2r* | Mm00439576_m1 | *RETRO Klf4** | FAM-CCCCTTCACCATGGCTG-MGB |
| *Kcnq1ot1* | Mm03936155_s1 | *RETRO Oct4** | FAM-CACCTTCCCCATGGCTG-MGB |
| *Kdr1* | Mm01222421_m1 | *Rian* | Mm01325842_g1 |
| *Klf4* | Mm00516104_m1 | *Slc38a4* | Mm00459056_m1 |
| *Magel2* | Mm00844026_s1 | *Sox2* | Mm03053810_s1 |
| *Mcts2* | Mm00481540_s1 | *Th* | Mm00447557_m1 |
| *Meox1* | Mm00440285_m1 | *Zdbf2* | Mm01254509_m1 |
| *Mest* | Mm00485003_m1 | *Zfp42* | Mm01194089 |
| *Nanog* | Mm02384862_g1 | *Zic1* | Mm00656094_m1 |
| *Nestin* | Mm00450205_m1 | | |

**Supplementary Table 4. List of Syber Green primers used.**

| ***Gene*** | **Forward (FW)** | **Reverse (RW)** |
|---|---|---|
| *Gnas* | AGAAGGACAAGCAGGTCTACCG | GTTAAACCCATTAACATGCAGGA |
| *Tsix* | TGTCAGGTTTCGGGGACACT | CTCTCCAGCCCAGGAACTGA |
| *Xist* | CTCATAGTAGTGGCCGACTA | TAAGCCCGTTAAGTAGTCCTT |

**Supplementary Table 5. List of pyrosequencing primers.**

| **DMR** | **Forward (FW)** | **Reverse (RW)** | **Sequencing** |
|---|---|---|---|
| *IG-DMR* | GTGGTTTGTTATGGGTAAGTTT | CCCTTCCCTCACTCCAAAAATTAA | GGTAAGTTTTATGGTTTATTGTATA |
| *Igf2r DMR1* | GGGATTTTAGAAAGATTGATTTTTTAAT | CCTCAAAACCAAACCTCAATTT | AATTTGGGTTTTTTTATTTAA |
| *Igf2r DMR2* | GGGTGAAGATTTTTGGGTTATAAG | CCCCCCCCAATACAACAA | TTTATTGTTTATTAGTGTTTTGAAT |
| *KvDMR* | AGAAGGGTGTTGAAGAAAAATT | ATCCTAAACCTAAACCTCCATAA | GTTGAGAAGTTAAGTGGA |
| *Mcts2* | TGAAGAAGAATTAGTGGGGTAA | ACAATTAAACACACTTTCCTTCTC | GGTGTTATTTTTTTTGTAGA |
| *Peg10* | AATTTTGTTAAGTTTTTAGTGGTTAGAT | CACTTAAAAATACAAAACCAATCACTT | CACAATTCCATCAATAACT |
| *Snrpn* | TTGGTAGTTGTTTTTTGGTAGGAT | TCCACAAACCCAACTAACCTTC | GTGTAGTTATTGTTTGGGA |
| *Zrsr1* | ATGGTTAGGTTGAGAGTTTTGGAAGTTT | TCCCTCAACAACCACTCTTCATA | TTTTGGAAGTTTTATTAGAGG |
